# Secretion-based production of prolyl-hydroxylated human type III collagen in scalable Physcomitrella photobioreactors

**DOI:** 10.1101/2025.11.26.690659

**Authors:** Lennard L. Bohlender, Juliana Parsons, Antonia Mitgau, Sebastian N. W. Hoernstein, Giovanna Grigolon, Bernhard Henes, Eva L. Decker, Ralf Reski

## Abstract

Collagens are structural proteins of the extracellular matrix essential for skin elasticity and integrity. They are widely used in dietary supplements and cosmetics. Conventional collagens of animal origin raise concerns regarding ethics, safety, and sustainability. As a vegan alternative, we report on the production of a 30 kDa prolyl-hydroxylated human collagen polypeptide from Physcomitrella moss plants. For secretion-based production and formulation compatibility, a hydrophilic region encompassing 334 amino acids from human type III collagen was selected, which includes four protein domains involved in cell adhesion, collagen binding, integrin recognition and wound healing. Transgenic moss lines were generated *via* protoplast transformation. Immunodetection identified collagen producing lines, and mass spectrometry validated the product and detected prolyl-hydroxylation on 23 sites. The presence of this important post-translational modification underscores the high biomimetic quality of the product. To enable industrial-scale production, the transformants were quantitatively analysed at the genomic, transcript, and protein levels. The most productive lines were forwarded to process development, where culture conditions including CO_2_ supplementation, pH, and light intensity were optimized. Upscaling to 5 L photobioreactors established a robust, light- and biomass-dependent production regime that yielded nearly 1 mg/L of secreted collagen polypeptide in the culture supernatant after 11 days of cultivation. Taken together, this study presents the first scalable moss-based production of a post-translationally modified human collagen and offers a sustainable and vegan alternative to conventional collagens for cosmetic formulations. This highlights the versatility of Physcomitrella as a production host for high-quality proteins with industrial applicability that also meet consumer requirements.

**Key message:** Scalable moss bioreactors enable the production of high-quality recombinant prolyl-hydroxylated human collagen without heterologous P4H expression, offering a sustainable and vegan alternative to conventional collagens derived from animals.

## Introduction

Human collagen is the dominant structural protein of the extracellular matrix (ECM) and the main component of connective tissue, where it contributes to its resilience and elasticity. Type III collagen provides tensile strength to hollow organs such as blood vessels, functions in cell adhesion, is involved in the blood-clotting cascade, and is important for wound healing (Ihlberg et al. 1993; Kim et al. 2005; Malfait 2018; Kuivaniemi and Tromp 2019).

Collagen is mainly obtained from animals such as cattle, pigs, rats, and fish, and it is widely used in biomedical materials, as an ingredient in dietary supplements and in cosmetics (Sionkowska et al. 2020). Due to growing market demands, safety concerns, including potential immunogenic reactions and the transmission of zoonoses, as well as consumer reservations about animal sources (Grigolon et al. 2023; Chen et al. 2024; Ariyanta et al. 2025), collagens are increasingly being biotechnologically produced in various non-animal derived expression systems (Rodríguez et al. 2018; Zhao et al. 2023; Chen et al. 2024). The market value of recombinant humanized collagen was approximately USD 1.5 billion in 2023 and is expected to reach USD 3.6 billion by 2032 (DataIntelo 2024).

In selecting a suitable production host, it is essential that post-translational protein modifications (PTM), in particular prolyl 4-hydroxylation by prolyl 4-hydroxylases (P4H), are carried out efficiently. The primary structure of collagen is defined by its highly repetitive Gly-Xaa-Yaa motif, with glycine occupying every third position and proline residues frequently occurring at both the Xaa and Yaa positions. Prolines at the Yaa position are frequently hydroxylated (Brodsky and Ramshaw 1997; Engel and Bächinger 2005). This site-specific prolyl-hydroxylation is an important PTM in collagen that is essential for its structural integrity, thermal stability, and biological function (Myllyharju 2008; Taga et al. 2021). Further, some collagen type and tissue-specific far less prevalent PTMs of collagens include prolyl 3-hydroxylation, which is absent in human collagen type III (Weis et al. 2010; Pokidysheva et al. 2013; Gjaltema and Bank 2016), lysyl hydroxylation (Walker et al. 2005; Gjaltema and Bank 2016), and occasionally hydroxylysine O-glycosylation (Peng et al. 2025) or extracellular lysyl oxidation (Kagan and Li 2002). These modifications cannot be fully reproduced in non-mammalian production hosts due to the absence of the required enzymes or appropriate substrate specificities.

Green systems used for the recombinant production of human collagen include the vascular plants barley, corn and tobacco (Shoseyov et al. 2014; Wang et al. 2017; Zhao et al. 2023). When produced in barley cell culture (Ritala et al. 2008) and barley seeds (Eskelin et al. 2009), corn grains (Zhang et al. 2009a; Zhang et al. 2009b) or tobacco (Ruggiero et al. 2000), the products showed little or no prolyl-hydroxylation and no additional detectable PTMs. Therefore, co-expression with different heterologous, non-plant P4Hs was achieved in tobacco plants (Merle et al. 2002; Stein et al. 2009) and corn grains (Xu et al. 2011) with higher yields of prolyl 4-hydroxylation.

The moss Physcomitrella (new botanical name: *Physcomitrium patens*) is a model plant for basic biology and biotechnology (Decker and Reski 2020; Lueth and Reski 2023) with a proven track record as a production host for biopharmaceuticals (Reski et al. 2015; Reski et al. 2018), including difficult-to-express proteins such as human factor H and its derivatives (Michelfelder et al. 2017; Top et al. 2019; Ruiz-Molina et al. 2022a), human erythropoietin (Weise et al. 2007; Parsons et al. 2012), spider silk (Ramezaniaghdam et al. 2025), and virus-like nanoparticles (Niederau et al. 2025). Unlike vascular plants, Physcomitrella can be grown in bioreactors (Hohe et al. 2002; Hiss et al. 2014) that can be operated in accordance with the principles of good manufacturing practice (GMP; Decker and Reski 2007). Moreover, its ability to secrete recombinant products directly into the simple mineral culture medium allows simplified downstream processing (Baur et al. 2005; Schaaf et al. 2005; Gitzinger et al. 2009), eliminating the need for tissue extraction and laborious purification steps. Furthermore, Physcomitrella is ideal for precise genome engineering due to efficient homologous recombination in mitotic cells (Strepp et al. 1998; Wiedemann et al. 2018), and its P4H enzymes are well characterised (Parsons et al. 2013; Rempfer et al. 2024). Non-transgenic mosses are rich in secondary metabolites (Erxleben et al. 2012; Horn et al. 2021; Munoz et al. 2024) and, when grown in photobioreactors, serve as a source of bioactive substances for the cosmetics industry, with the first products already on the market (Wandrey et al. 2018). Therefore, we decided to investigate whether a prolyl-hydroxylated human collagen polypeptide could be produced in Physcomitrella without the co-expression of heterologous P4H enzymes and whether it would be secreted into the simple mineral medium, which would distinguish it from recombinant collagens produced in other plant systems.

Here, we identified a fragment of 334 amino acids (aa) from the human collagen α1(III) chain (P02461, COL3A1) that comprises four protein domains important for bioactivity and is hydrophilic. Subsequently, we optimized the process with a focus on three major objectives: construct design, validation of post-translational modifications, and a scalable secretion process. The collagen-coding sequence was optimized for expression in Physcomitrella without altering the aa sequence of human collagen. The transgenic moss lines were examined quantitatively at the genomic, transcript and protein levels. The product was secreted into a simple mineral medium, harvested and characterized. Analysis by mass spectrometry provided evidence for an intact product with prolyl-hydroxylation. A bioreactor process was established and optimized, resulting in up to 1 mg/L of secreted prolyl-hydroxylated human collagen polypeptide in the culture supernatant after 11 days of cultivation.

## Results

### Polypeptide selection and codon optimization

In this study, we aimed to establish the recombinant production of a vegan collagen alternative to animal-derived collagens for use in cosmetic formulations, employing the moss Physcomitrella as a production platform. Given its crucial role in wound healing (Krafts 2010; Stewart et al. 2025), skin integrity, and tissue regeneration (Volk et al. 2011; D’hondt et al. 2018), human collagen type III was selected as the basis for the recombinant polypeptide. With regard to compatibility with a secretion-based production system and broad applicability in cosmetic formulations, particular attention was paid to selecting a predominantly polar protein sequence.

Therefore, the full-length sequence of human pre-procollagen α1(III) (1466 aa, including the signal peptide; UniProt ID: P02461), from which the collagen triple helix-forming collagen α1(III) chain is formed after cleavage of the N- and C-terminal propeptides, was examined *in silico* for its hydropathicity. This analysis identified an extended hydrophilic region of 334 aa within the collagen α1(III) chain extending from glycine 456 to glycine 789 (**Figure 1**). With respect to hydroxylation, this region includes 49 4-hydroxylated prolines, no 3-hydroxylated prolines, and no hydroxylated lysins in humans (Kuivaniemi and Tromp 2019). In addition, this polypeptide sequence contains four protein domains that are bioactive and are positively associated with skin integrity: I) The domain Gly489–Gly510, which is involved in collagen binding (Hua et al. 2019), II+III) Two integrin-binding motifs (GAOGER, GMOGER; O: hydroxyproline; Siljander et al. 2004; Kim et al. 2005), and IV) A substantial portion of the Gly426–Ser560 region, which can increase type I collagen production and promote wound healing *in vitro* (Kim et al. 2024) (**Figure 1**). Our focus was on solubility of the product and not necessarily on triple helix formation. The resulting polypeptide has a calculated molecular weight of 30 kDa and an isoelectric point (pI) of 6.65 (ExPASy Compute pI/Mw; https://web.expasy.org/compute_pi/).

**Fig. 1.**
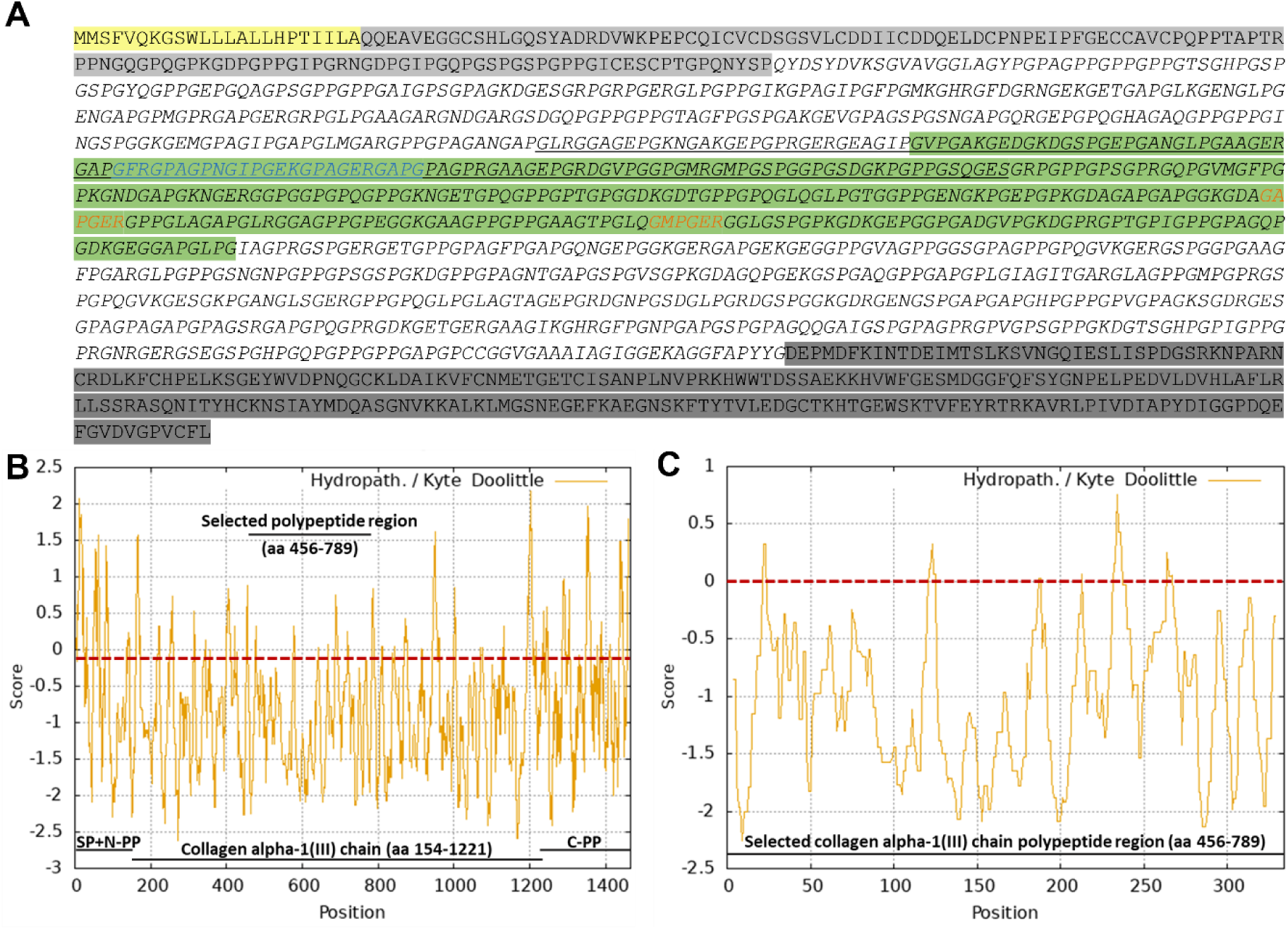
Full-length amino acid sequence of the human pre-procollagen α1(III) and corresponding hydropathicity plots. **A** Full-length amino acid sequence of the human pre-procollagen α1(III) (P02461), consisting of the signal peptide (yellow; aa 1–23), the N-terminal propeptide (light grey; aa 24–153), the collagen α1(III) chain (italics; aa 154–1221), and the C-terminal propeptide (dark grey; aa 1222–1466). The sequence region selected for recombinant production is highlighted in green (aa 456–789), integrin recognition motifs are marked in orange, the segment Gly426-Ser560 involved in collagen-related signalling and interactions is underlined, and the collagen binding Gly489-Gly510 sequence stretch within this region is emphasized in blue. **B** *In silico* hydropathicity plot of the human pre-procollagen α1(III) aa sequence. **C** *In silico* hydropathicity plot of the selected 334-aa region. Hydropathicity plots were generated using the ExPASy ProtScale tool (https://web.expasy.org/protscale/) with the Kyte and Doolittle (1982) scale. Positive values indicate hydrophobic amino acids, and negative values indicate hydrophilic amino acids. Abbreviations: SP: signal peptide; N-PP: N-terminal propeptide; C-PP: C-terminal propeptide; aa: amino acid.

The coding sequence (CDS) of the selected COL3A1 polypeptide underwent three rounds of *in-silico* optimization. First, the codons were adapted towards the Physcomitrella codon usage. Subsequently, nucleotides within 11 heterosplicing recognition motifs (AGGT, Top et al. 2021) were altered. In a final step, eight codons that are underrepresented in Physcomitrella (Nakamura et al. 2000; Hiss et al. 2017), were replaced with the corresponding overrepresented alternatives (**Supplementary Table S1**) using the online tool physCO (Top et al. 2021). This resulted in 192 base changes at the nucleic acid sequence level (**Supplementary Figure S1**) but did not alter the resulting aa sequence of the human collagen polypeptide. After *in-silico* fusion of the CDS to the PpAP1 (Pp6c5_10120V6.1) signal-peptide (Schaaf et al. 2004) to enable product secretion and the addition an 8x His-tag, the synthesized CDS (GeneArt, Thermo Fisher Scientific) was cloned into an Actin5 promoter-driven (Weise et al. 2006; Niederau et al. 2024) expression vector, which additionally contained an hpt cassette (Decker et al. 2015) for selection purposes (**Supplementary Figure S2**).

### Immunodetection and mass spectrometric sequence analysis

Following PEG-mediated transformation of Physcomitrella wild type (WT) protoplasts, their regeneration and selection, 50 hygromycin-resistant moss lines were cultivated in agitated flasks and examined for collagen production using Western blot analysis. For this purpose, total soluble proteins were extracted from protonema, the young filamentous tissue of the moss, which was cultivated under standard conditions. The collagen polypeptide was enriched using ion affinity chromatography (His SpinTrap) and analysed by Western blot using anti-His antibodies. Collagen polypeptide-producing lines showed a strong signal at approximately 45 kDa. In addition, an unspecific band at around 37 kDa was consistently detected in all analysed lines, including WT. The eight lines with the strongest signals (C6.1, C5, C11, C13, C15, C34, C44, and C46) are shown alongside a WT control in **Figure 2A**. To verify that the 45 kDa signal originated from the recombinantly produced collagen polypeptide, the purified extract from line C6.1, which together with C5 showed the strongest anti-His antibody signals, was further analysed using an anti-collagen antibody-based immunodetection. The resulting signals were consistent with those obtained using the anti-His antibody (**Figure 2B**), confirming that the bands detected at approximately 45 kDa correspond to the recombinant collagen polypeptide. To analyse the quality of the recombinant product, the protein band corresponding to the 45 kDa signal was excised from an SDS-PAGE gel prepared and run in parallel (**Figure 2B**) and analysed by mass spectrometry. This analysis confirmed the successful production of the intact full-length collagen polypeptide and confirmed the proper cleavage of the signal peptide (**Figure 2C**). Of the eight lines compared, five lines (C5, C6.1, C15, C34, and C44) were selected for further analysis.

**Fig. 2.**
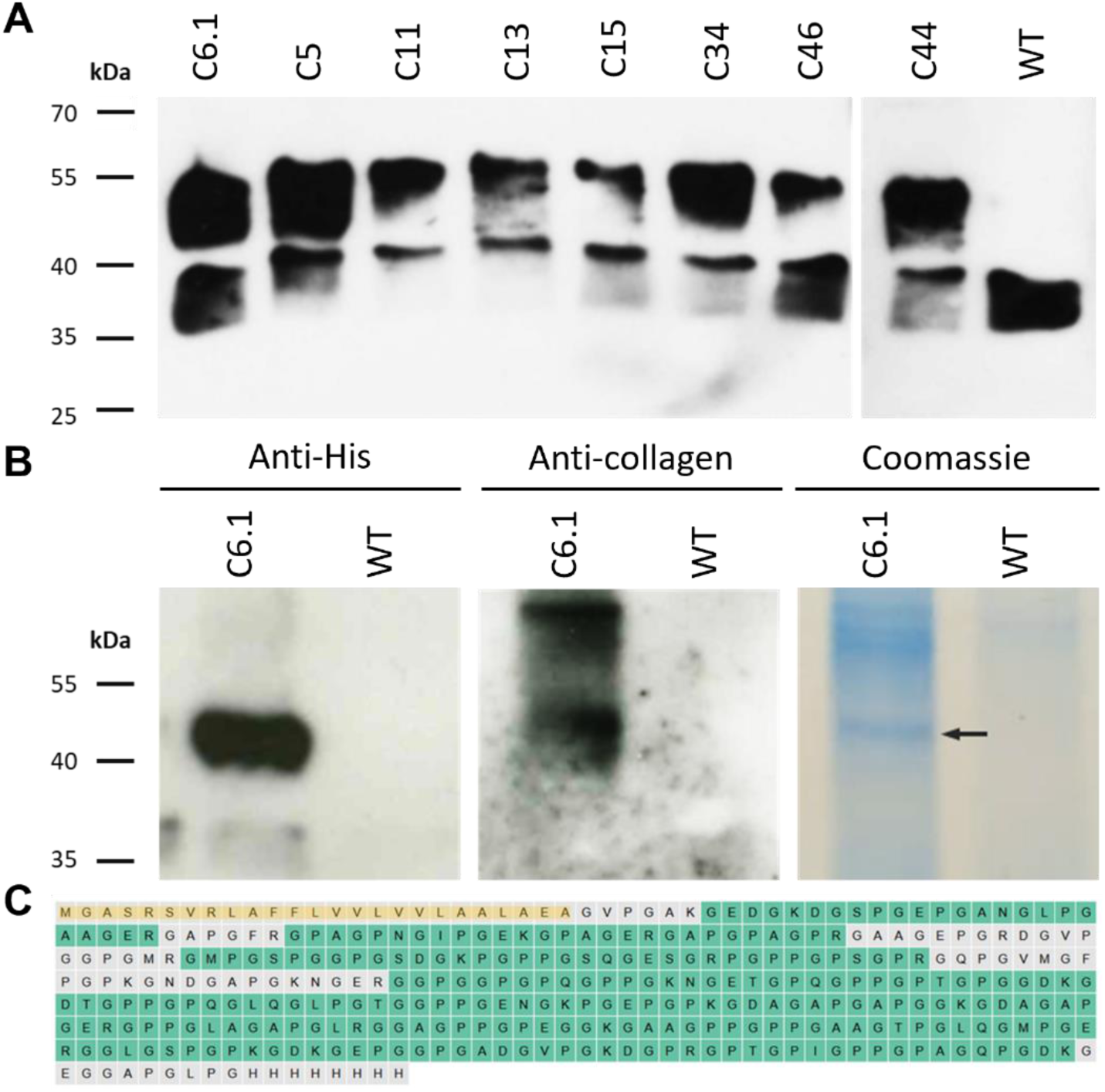
Analysis of the Physcomitrella-produced recombinant collagen polypeptide. **A** Western blot using anti-His antibodies of 10 µg His SpinTrap-enriched proteins from protonema tissue of transgenic Physcomitrella lines producing recombinant collagen polypeptide, compared with a WT control. **B** Co-localization of signals confirmed by anti-His and anti-collagen immunodetections on His SpinTrap-enriched proteins from line C6.1. **C** Mass spectrometric analysis of the intracellularly accumulated recombinant collagen polypeptide revealed 81% sequence coverage (identified aa are shown in green, non-identified in grey) of the mature polypeptide and confirmed cleavage of the signal peptide (highlighted in yellow). Uncropped immunodetection and SDS-PAGE images are provided in **Supplementary Figure S3**.

### Quantitative analysis at genomic, transcript and protein level

To compare the production yields of the five selected lines C5, C6.1, C15, C34 and C44, the produced collagen polypeptide was quantified using ELISA. Protonema suspension cultures of the selected lines, along with a WT control, were adjusted to a density of 60 mg dry weight (DW) per litre, cultivated under standard conditions and harvested after seven days of cultivation. Extracted total proteins were analysed using an anti-His antibody-based ELISA, which detects both monomeric forms and higher-order assemblies of the product. Signals were blank-corrected, WT-subtracted, and calibrated against a His-tagged elastin peptide standard curve. Lines C5 and C6.1 produced the highest amounts of collagen polypeptide, with approximately 30 µg and 32 µg per gram of protonema fresh weight (FW), respectively. Line C15 yielded an intermediate amount of around 19 µg/g FW, while the lowest levels were observed in lines C34 and C44, with approximately 8 µg/g FW and 9 µg/g FW (**Figure 3A**).

**Fig. 3.**
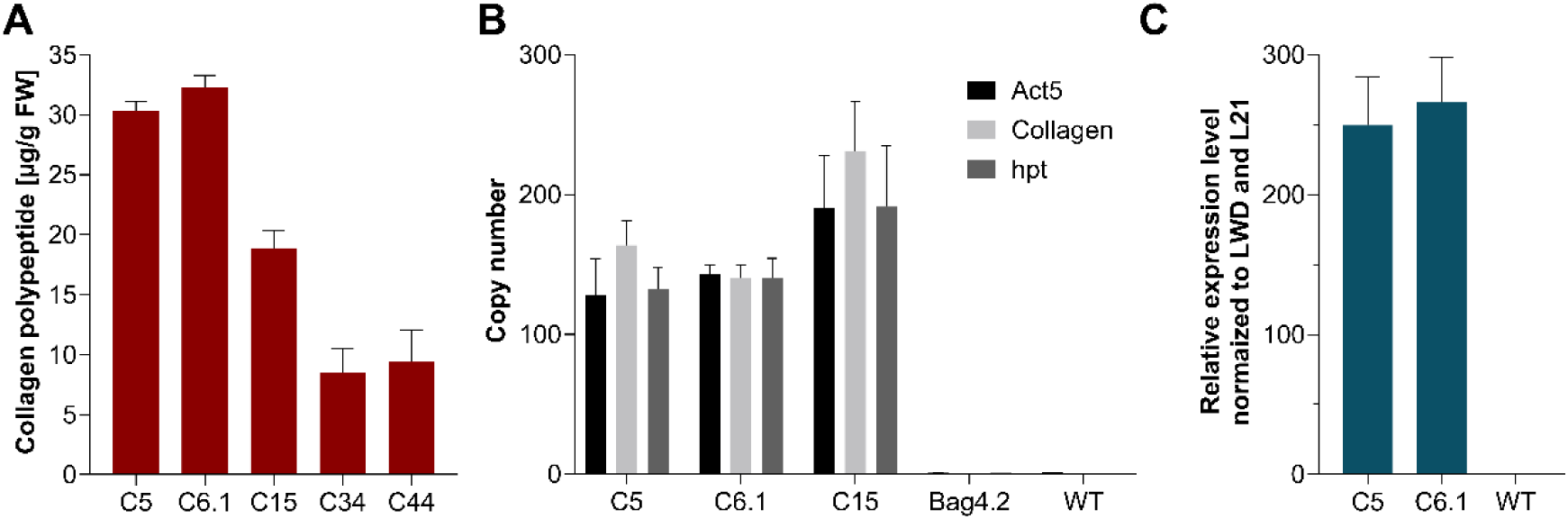
Quantitative analysis of intracellular yields, construct copy numbers, and expression levels in collagen-producing moss lines. **A** Intracellular accumulation of recombinant collagen polypeptide in producing lines, quantified by ELISA. Bars represent blank-corrected and WT-signal subtracted values (mean ± SD, n = 3 technical replicates). **B** qPCR analysis of construct copy numbers in the three best-producing lines using primer pairs targeting the Actin5 promoter (Act5), the collagen CDS (Collagen), and the hygromycin resistance cassette (hpt). Normalization was performed against signals from two primer pairs targeting the endogenous single-copy CLF gene. WT and the confirmed hpt single-integration line Bag4.2 served as references for normalization of Act5 and hpt-derived signals, respectively. Bars represent normalized mean values ± SD (n = 3 technical replicates). **C** qRT-PCR analysis of collagen expression levels in the two best-producing lines compared to WT. Expression was measured with a primer pair targeting collagen cDNA and normalized to the housekeeping genes LWD and L21. Relative expression is shown as 2^−ΔCT^, where ΔCT = CT(transgene) - CT(housekeeping). Bars represent mean ± SD (n = 3 technical replicates).

For molecular characterization at the genomic level, the number of construct copies that had been integrated into the moss genome was determined by quantitative PCR (qPCR). This analysis was performed on the three best-producing lines using primer pairs targeting the Actin5 promoter, the collagen CDS, and the hpt cassette, respectively. In addition, all five selected lines were compared using only the Actin5 promoter- and collagen-specific primers. Alongside the collagen-producing lines, the WT and line Bag4.2, a known single-integration line for the hpt CDS (Bohlender et al. 2022), were included as references for relative quantification of Actin5- and hpt-derived signals. To normalize for genomic DNA input, integration signal intensities were obtained for each line using two primer pairs targeting the CLF gene, a confirmed single-copy gene in Physcomitrella (Pereman et al. 2016). The qPCR analysis revealed that line C15 carried the highest construct copy number, with approximately 200 integrations. Lines C5 and C6.1 followed, each harbouring around 140 integrations (**Figure 3B**). In contrast, lines C34 and C44 contained lower copy numbers, with about 20 and 30 integrations, respectively (**Supplementary Figure S4**). Thus, it became evident that the highest copy number does not guarantee the highest product yield, reinforcing an important practical lesson for transgenic production systems.

To compare the two best-producing lines C5 and C6.1 at the transcript level, collagen CDS expression levels were quantified by qRT-PCR. Total RNA was extracted from protonema material of suspension cultures initiated with 60 mg DW/L and harvested seven days after inoculation, then reversely transcribed into cDNA. Expression levels were normalized to the two strongly expressed housekeeping genes LWD (Pp6c22_10850V6.1; Yin et al. 2021) and L21 (Pp6c13_1120V6.1; Beike et al. 2015). This analysis showed that line C5 exhibited an approximately 250-fold, and line C6.1 a 266-fold higher expression of the collagen CDS relative to the two housekeeping genes (**Figure 3C**). Based on these quantitative analyses, the two best performing collagen-producing lines C5 and C6.1 were chosen for the establishment of a secretion-based production process.

### Optimization of production parameters

To establish a secretion-based production process suitable for the downstream industrial formulation of the recombinant collagen polypeptide, cultivation parameters influencing the secretion and accumulation of recombinant human collagen in the culture medium were investigated and optimized. For this purpose, 500 mL agitated Erlenmeyer flasks were inoculated with 200 mL of protonema suspension culture from either line C5 or the WT, adjusted to a density of 300 mg DW/L. Cultures were grown for 16 days under standard conditions (Decker et al. 2015) or with supplementation of 2% CO_2_ in the incubator atmosphere (20,000 ppm CO_2_). Supernatant samples were collected on days 3, 6, 9, 14, and 16 and analysed by ELISA. In both conditions, the highest collagen polypeptide accumulation occurred on day 6, followed by a decline in supernatant concentrations toward the end of cultivation. Under 2% CO_2_ supplementation, approximately 50 µg/L collagen polypeptide was achieved, more than twice the amount compared to the ∼22 µg/L achieved under standard atmospheric conditions (**Figure 4A**).

**Fig. 4.**
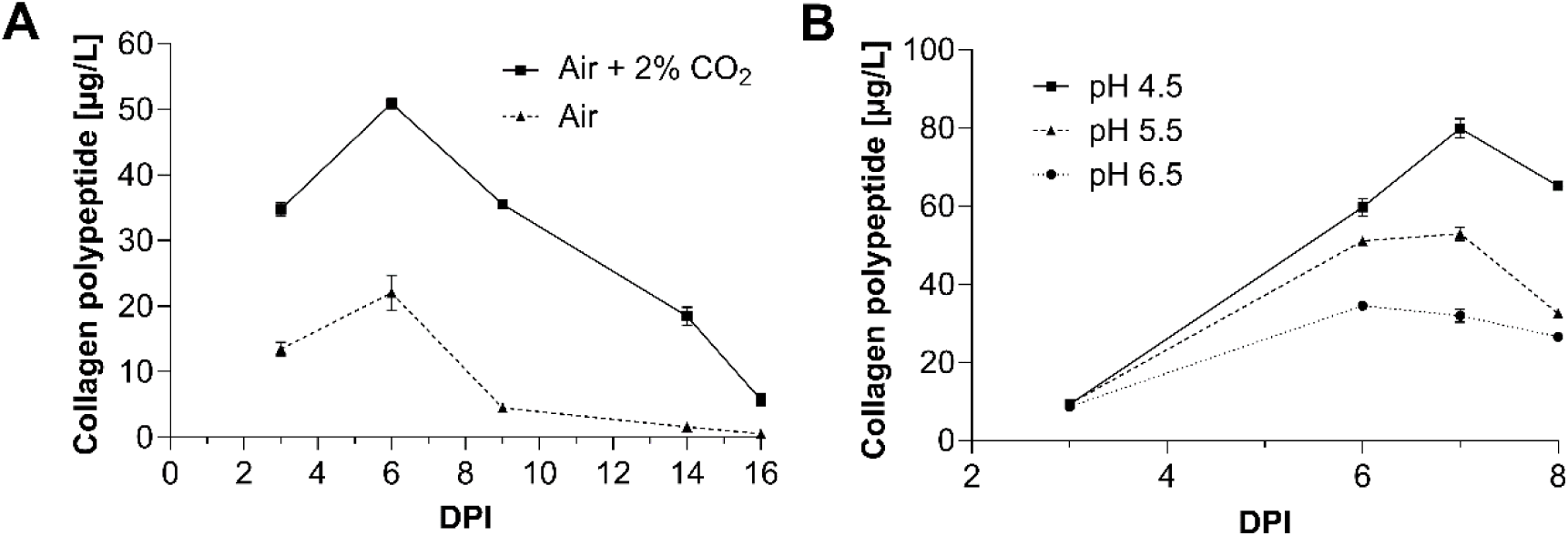
Effects of cultivation conditions on recombinant collagen polypeptide accumulation in the culture supernatant. Secreted collagen polypeptide levels from cultures inoculated with protonema tissue (300 mg DW/L) of line C5 and WT controls were quantified by ELISA. **A** Cultivation for 16 days under standard conditions (Air) or with 2% CO_2_ supplementation (Air + 2% CO_2_). **B** Cultivation for 8 days at pH 4.5, 5.5, or 6.5 with 2% CO_2_ supplementation. Data points represent blank-corrected, WT-subtracted mean values ± SD (n = 3 technical replicates). DPI: day(s) post inoculation.

Next, the effect of medium pH on collagen polypeptide accumulation was examined under CO_2_ supplementation. The medium was adjusted to pH 4.5, 5.5, or 6.5, and product levels were monitored over 8 days. The highest accumulation was observed in medium adjusted to pH 4.5, which reached about 80 µg/L on day 7. At pH 5.5, accumulation peaked at about 55 µg/L on the same day, while at pH 6.5 about 37 µg/L at its maximum on day 6 were achieved (**Figure 4B**). Here, we focussed on quantitative yield and did not monitor protease activity.

Collectively, these optimization experiments identified an increased CO_2_ availability of 2% and medium pH adjusted to 4.5 as the most favourable conditions for collagen polypeptide accumulation in the culture supernatant. These optimized conditions were subsequently applied in 5 L photobioreactor runs for scale-up and industrial formulation.

### Upscaling to 5 L photobioreactors

To provide the recombinant collagen polypeptide in amounts suitable for industrial formulation, the secretion-based production process was scaled up to 5 L photobioreactors. Therefore, based on the information gained from our previous results, a 5 L photobioreactor was set up with a pH controlled at 4.8 and aerated with air enriched with 2% CO_2_. The bioreactor was inoculated with line C5 at an initial biomass density of 400 mg DW/L and maintained for 22 days. Light intensity was set to 90 µmol m^-2^ s^-1^ until day 9, then set to 190 µmol m^-2^ s^-1^ until day 14, followed by 380 µmol m^-2^ s^-1^ until day 17, and finally set to 760 µmol m^-2^ s^-1^ until the end of the bioreactor run. Biomass growth was monitored, and supernatant as well as biomass samples were collected for further analysis.

Over the course of the bioreactor run, secreted collagen polypeptide concentrations increased first slowly from 3 µg/L on day 3 to 250 µg/L on day 17 and then strongly reaching a maximum of 986 µg/L on day 21, before slightly declining on day 22 (**Figure 5A**). Intracellular collagen levels remained low throughout, peaking at 7 µg/g fresh weight on day 7 (**Figure 5A**). The biomass initially increased almost linearly from 400 mg DW/L to 1800 mg DW/L by day 7, then gradually reached approximately 2700 mg DW/L by day 17, followed by a stronger increase to 4200 mg DW/L until day 22 (**Figure 5A**). This initial process yielded nearly 1 mg of collagen polypeptide per litre of culture supernatant within a production period of 22 days.

**Fig. 5.**
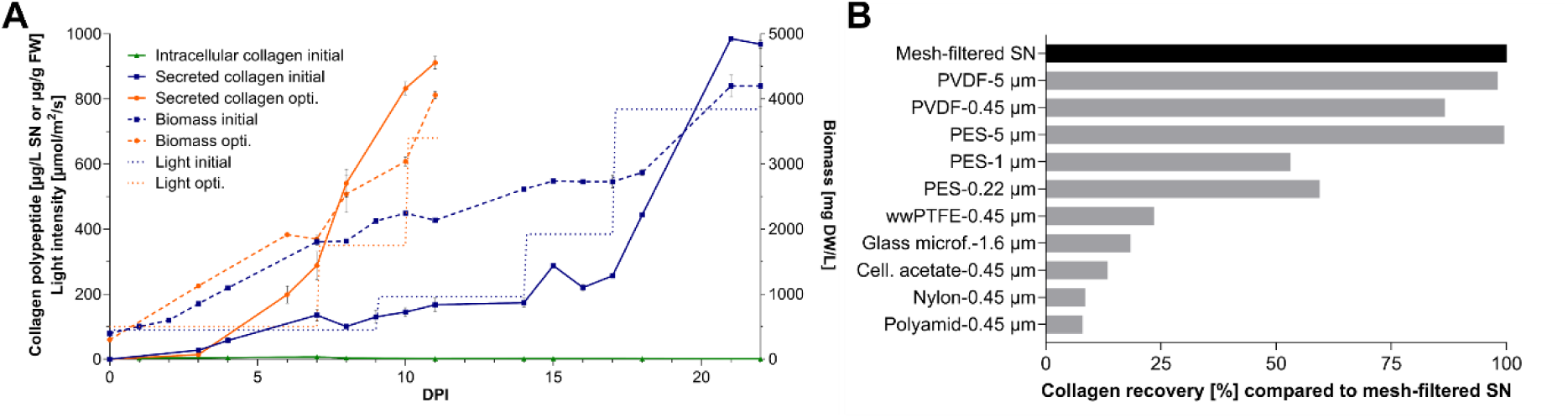
Bioreactor culture parameters and clarification of recombinant collagen polypeptide containing supernatant. **A** Collagen polypeptide accumulation profiles (solid lines), moss biomass accumulation (dashed lines), and applied light intensity settings (dotted lines) during 5 L photobioreactor runs of line C5 under initial and optimized conditions. Collagen concentrations were quantified by ELISA (mean ± SD, technical triplicates) and the biomass was analysed by dry weight measurements (mean ± SD, technical duplicates). For improved readability, a split version of this panel is provided in **Supplementary Figure S5**. **B** Recovery efficiency of the collagen polypeptide (in %) after an additional filtration step using different filter materials, relative to the 11 µm mesh-filtered supernatant. Abbreviations: opti.: optimized; SN: supernatant, FW: fresh weight, DW: dry weight, DPI: day(s) post inoculation, Glass microf.: Glass microfiber, Cell. acetate: Cellulose acetate.

To develop a more time- and cost-efficient production process, conclusions were drawn from several bioreactor runs conducted under different parameter regimes. Evaluation of these data led to the establishment of a biomass-dependent, light-intensity-regulated production process. A light intensity of 100 µmol m^-2^ s^-1^ was applied up to a biomass of 2 g DW/L, followed by a light intensity of 350 µmol m^-2^ s^-1^ until a biomass density of 2.8 g DW/L, which was then increased to 680 µmol m^-2^ s^-1^ until the harvest of the supernatant, which was performed at a biomass density of about 4 g DW/L. It appeared that biomass accumulation higher than 4 g DW/L resulted in a decrease of the amount of collagen polypeptide in the culture supernatant. With the optimized production conditions, product concentrations exceeding 900 µg/L were achieved after approximately 11 days of cultivation (**Figure 5A**). This effectively halved the original production time while maintaining comparably high product yields. This production regime is likely to be product-specific and reinforces the need to optimize upstream processing for each new product.

To render the collagen polypeptide-containing supernatant directly suitable for downstream industrial formulation, a clarification protocol was established. First, the bioreactor material was harvested under sterile conditions, and the biomass was removed from the culture supernatant through two sequential filtration steps using 100 µm and 11 µm mesh polypropylene sieves. To ensure a product free of residual cells and debris, the suitability of the following filter materials was assessed for a subsequent filtration step: cellulose acetate (0.45 µm), glass microfiber (1.6 µm), nylon (0.45 µm), polyethersulfone (PES; 0.22, 1, and 5 µm), polyamide (0.45 µm), polyvinylidene difluoride (PVDF; 0.45 and 5 µm), and water-wettable polytetrafluoroethylene (wwPTFE; 0.45 µm). Collagen polypeptide concentrations in these filtrates and in the 11 µm mesh-filtered control were quantified by ELISA, and recoveries were calculated as percentages of the initial product concentration present prior to the additional filtration step. When cellulose acetate, glass microfiber, nylon, polyamide and wwPTFE filters were used, less than 25% of the initial product was recovered. In contrast, filtration with PVDF and PES filters preserved nearly 100% of the initial product. This was particularly evident with filters of 5 µm pore size, where almost no collagen loss was detected. At smaller pore sizes, 87% of the initial collagen concentration was recovered with 0.45 µm PVDF filters, whereas 1 µm and 0.22 µm PES filters resulted in a recovery of 50-60% (**Figure 5B**).

Taking all results into consideration, PVDF filter material with a pore size of 0.45 µm provided the best relationship between collagen recovery and filtration effect, and thus represents a promising option for integration into the production process.

### Evaluation of product quality

Following the establishment of the secretion-based production process, the quality of the product in the culture supernatant was evaluated using Western blot-based immunodetection and mass spectrometry.

The immunodetection of precipitated supernatants from C6.1 and WT bioreactor runs revealed a band at approximately 37 kDa in the supernatant of line C6.1, along with a very intense high molecular weight signal ranging from 45 to over 130 kDa, while no signals were detected in the sample of the WT supernatant (**Figure 6A**). To check if the detected signals originated from the collagen polypeptide, the corresponding regions were excised from an SDS-PAGE gel run in parallel and analysed by mass spectrometry. These analyses confirmed the presence of the collagen polypeptide in all signal-associated gel regions, verified the correct cleavage of the AP1 signal peptide, and yielded a combined sequence coverage of 58% of the mature collagen polypeptide (**Figure 6B**). In addition, these measurements confirmed the presence of hydroxyproline residues (**Figures 6B, C**) as a consequence of enzymatic prolyl-hydroxylation (Rempfer et al. 2024). In the secreted collagen fraction, a total of 41 proline residues were identified. Of these, 28 were located at the Yaa position and 13 at the Xaa position within the characteristic collagen Gly-Xaa-Yaa amino acid motif. Among all identified proline sites, 23 (∼54%) were hydroxylated, with site-specific occupancy ranging on average from 10% to 80% (**Figure 6C**). Seventeen (∼74%) of these hydroxylated prolines were found at Yaa positions, including the hydroxylated GAOGER integrin-binding motif (**Figure 6B**) and the remaining six occurred at Xaa positions. Importantly, mass spectrometric analyses provided no evidence for plant-specific *O*-glycosylation of hydroxyprolines, such as the attachment of up to five arabinose residues or single hexoses. Consistent with these results, Western blot analysis with the anti-1,5-α-L-arabinan antibody LM6-M showed no evidence for arabinogalactosylation of the secreted collagen peptide (**Supplementary Figure S6**).

**Fig. 6.**
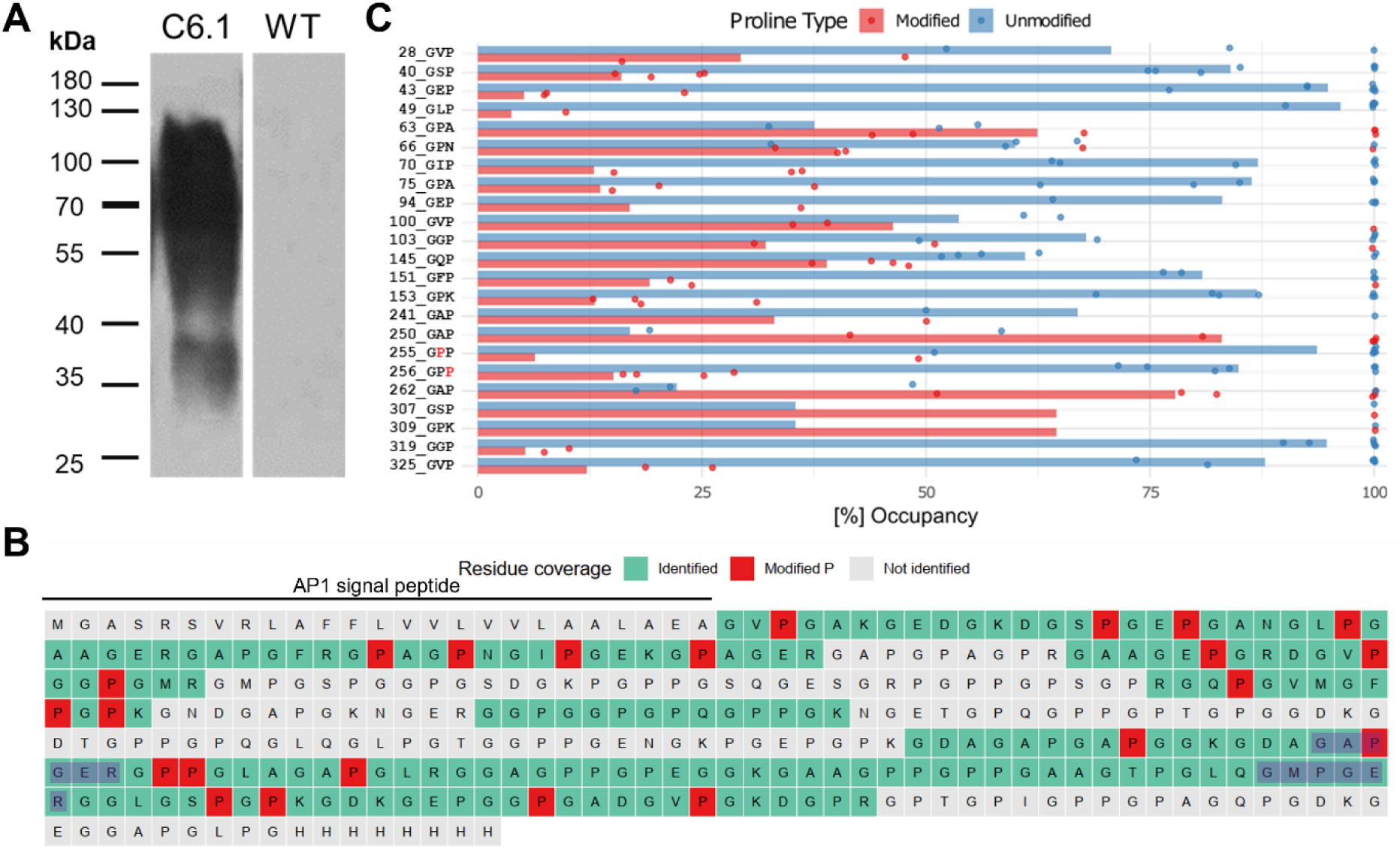
Evaluation of the secreted recombinant collagen polypeptide in bioreactor supernatants. **A** Anti-His immunodetection of acetone-precipitated bioreactor supernatants (1.5 mL) from a bioreactor run of line C6.1 compared with an equally treated WT control. Mass spectrometry confirmed the presence of the recombinant collagen polypeptide in all detected signal areas (**Supplementary Figure S7**). **B** Combined sequence coverage of the secreted collagen polypeptide obtained from seven mass spectrometric measurements of the bioreactor supernatant. Identified sequence regions are highlighted in green, hydroxylated proline residues in red, and non-identified regions in grey. The length of the secretion-mediating AP1 signal peptide is indicated above and the two sequence regions of the two integrin binding motifs GAOGER and GMOGER (O: hydroxyproline) are highlighted in blue. **C** Chart shows the percentage of hydroxylation of prolines found at each of the 23 modified positions identified over seven measurements of secreted collagen polypeptide containing bioreactor supernatant samples. Sites within the G-X-Y motif are shown in order of their occurrence along the amino acid sequence. Only prolines with a PTMProphet site localization probability > 0.75 were used. As quantitative values, label-free quantitation intensities were used, log2 transformed and normalized to the median intensity of each measurement. Subsequently, all values were centred to the median across all measurements. Data points represent percentages related to the single experiments whereas bars represent percentages across all measurements. A list of all measured hydroxyproline sites and their intensities is available in **Supplementary Table S3**.

Finally, Western blot analysis of the non-precipitated supernatant revealed that under the applied conditions the majority of the product accumulated in a molecular weight range between 70 and 100 kDa (**Supplementary Figure S7**).

At the current stage, the established 5 L photobioreactor process, combined with sequential filtration through 100 µm and 11 µm mesh polypropylene sieves, yields a final product containing nearly 1 mg/L recombinant collagen polypeptide. Following sterile harvesting, the product will be subsequently subjected to further filtration, *in-vitro* activity testing, and formulation.

## Discussion

Collagens have gained increasing popularity as cosmetic ingredients in recent years. However, the vast majority of available products still rely on animal-derived sources (Sionkowska et al. 2020). Animal collagens pose potential risks, including contamination with human pathogens and may trigger immune reactions (Mullins et al. 1996; Mitura et al. 2020; Ariyanta et al. 2025). Recombinant collagen production offers clear advantages, including high purity, low immunogenicity, and alignment with the growing consumer demand for vegan, sustainable, and ethically sourced ingredients (Chen et al. 2024). To address these demands, this study describes the production of a prolyl-hydroxylated recombinant human type III collagen polypeptide in transgenic Physcomitrella moss plants, alongside the development of a secretion-based production process in 5 L photobioreactors.

Among the 28 known collagen types (Gordon and Hahn 2010; Ricard-Blum 2011), human type III collagen was selected for recombinant production due to its critical roles in wound healing (Krafts 2010; Stewart et al. 2025), tissue formation, fibrillogenesis (Liu et al. 1997; Kuivaniemi and Tromp 2019), and the maintenance of skin integrity and regeneration (Volk et al. 2011; D’hondt et al. 2018).

Within the human collagen type III sequence, a 334 aa long segment (Gly456 - Gly789) was computationally identified as a promising candidate for recombinant production. This region, representing 31% of the triple-helix-forming human collagen α1(III) chain, shows low hydrophobicity, favourable aqueous solubility, and contains four domains implicated in the modulation of the extracellular matrix (ECM), and thus bioactivity. These features make it particularly well-suited for secretion and formulation purposes, defining it as a compelling candidate for recombinant production as a bioactive cosmetic ingredient.

Codon-optimized collagen-encoding expression constructs were introduced in Physcomitrella protoplasts by PEG-mediated protoplast transformation. Immunodetection of purified total protein extracts of producing lines revealed specific bands at approximately 45 kDa, exceeding the predicted molecular weight of the collagen polypeptide of 30 kDa. This shift is consistent with the extended conformation of collagens in SDS-PAGE, a known phenomenon for both recombinant collagen products and commercial collagen standard proteins (Aly et al. 2022).

To enable the production of the recombinant collagen polypeptide at an industrially relevant scale, successful Physcomitrella lines were identified by quantifying construct copy numbers *via* qPCR and intracellular collagen accumulation *via* ELISA. In four of the five tested lines, accumulation levels positively correlated with construct copy numbers, consistent with previous observations in transgenic Physcomitrella lines (Ruiz-Molina et al. 2022b; Saliba et al. 2026). The highest intracellular collagen polypeptide levels, with 30 µg/g FW and 32 µg/g FW, respectively, were obtained from lines C5 and C6.1, both carrying approximately 140 integrated construct copies. In contrast, line C34, with about 20 integrations, produced 8 µg/g FW, representing the lowest yield among the tested lines. Interestingly, line C15, despite harbouring ∼200 copies, accumulated only 19 µg/g FW, possibly due to construct insertions disrupting endogenous genes. Because lines C5 and C6.1 showed nearly identical copy numbers, yields, and expression levels, both were considered suitable for establishing a scalable production pipeline.

Secretion of the collagen polypeptide into the culture medium, enabled by a signal sequence (Schaaf et al. 2005) and by its deliberate hydrophilicity, was confirmed and optimized in small-scale flask experiments. The highest extracellular collagen accumulation was achieved under 2% CO_2_ supplementation. A medium pH of 4.5 outperformed the accumulation yields achieved at pH levels of 5.5 and 6.5, respectively. This is consistent with the calculated isoelectric point of the collagen polypeptide (pI = 6.65). Near the pI, peptides exhibit minimal net charge and tend to aggregate (Vate et al. 2023), while at pH < pI they carry a net positive charge, enhancing their water solubility (Wu et al. 2024). Thus, the increased yields at acidic pH reflect a greater solubility and stability under these conditions. In addition, alterations in pH may alter the activity of proteases in the culture medium.

The optimized parameters were successfully scaled up to 5 L photobioreactors, and a robust light- and biomass-oriented production process was developed. Over 11 days of cultivation, this process yielded nearly 1 mg collagen polypeptide per litre of culture supernatant, providing a suitable product for further industrial formulation. However, further strain engineering, including the optimization of promoters and terminators for heterologous expression (Horstmann et al. 2004; Gitzinger et al. 2009; Niederau et al. 2024), the addition of stabilizing additives (Baur et al. 2005), and a further upscaling to 500 L bioreactors (Michelfelder et al. 2017) can improve the yield of the product.

While Western blot and mass spectrometry analyses of the intracellularly accumulated collagen polypeptide already confirmed the synthesis of the full-length product with a nearly complete sequence coverage accumulating in a single band at a molecular weight of about 45 kDa, the quality of the secreted product from photobioreactors was also examined. Immunodetection of precipitated bioreactor supernatants revealed a band at ∼37 kDa corresponding to the collagen polypeptide monomer alongside intense higher molecular weight signals ranging from 45 to over 130 kDa. Mass spectrometric analyses confirmed the presence of the collagen polypeptide in all signal-related areas and confirmed the correct cleavage of the AP1 secretion signal peptide (Schaaf et al. 2004). The slight size difference between the monomeric intra-and extracellular product likely reflects differences in the sample matrices, as buffer components and co-purified proteins can influence peptide migration in the SDS-PAGE (See et al. 1985; Rath et al. 2009). According to MS analysis and Western blot analysis these differences in migration patterns are not due to hydroxyproline-*O*-glycosylation, which adds arabinoses or arabinogalactan to the protein (**Supplementary Figure S6**). This modification is common in vascular plants and mosses (Lee et al. 2005; Bohlender et al. 2022; Stenitzer et al. 2022), but did not occur here in the recombinant human collagen polypeptide.

Several other factors could have influenced the migration pattern in the SDS-PAGE. The signals observed with higher molecular weight present in the secreted fraction may possibly be due to the cross-linking of collagen polypeptides, a known property of collagens that contributes to tissue stability (Tanzer 1973). While this process is mainly catalysed by lysyl oxidase, non-enzymatic spontaneous crosslinking has also been reported (Gaar et al. 2020). Such spontaneous *in vitro* dimerization and multimerization may explain some of the higher molecular weight signals detected by immunoblotting. In addition, collagens are capable of forming both covalent and non-covalent crosslinks with polyphenols. Non-covalent interactions are promoted at acidic pH values (4-5), where hydrogen bonding and electrostatic interactions between protonated collagen amino groups and polyphenol hydroxyl groups are favoured (Wu et al. 2019). Such polyphenol crosslinks can enhance collagen biostability and protect against proteolytic degradation (Madhan et al. 2005; Shavandi et al. 2018; Wu et al. 2019; Hass et al. 2021). Since the presence of polyphenols in Physcomitrella cultures is well documented (Richter et al. 2012; Renault et al. 2017; Kriegshauser et al. 2021), such interactions could potentially be responsible for three observed phenomena: I) the increased product accumulation at pH 4.5 compared to higher pH values (5.5 and 6.5), II) the occurrence of additional high molecular weight signals in precipitated supernatant samples, and III) the lower sequence coverage achieved for the secreted collagen polypeptide compared to the intracellular collagen polypeptide. In the latter case, cross-linking could mask peptides, making it difficult to identify them using mass spectrometry.

Additionally, immunodetection of the non-precipitated bioreactor culture supernatant showed that under our conditions, the majority of the product accumulates between 70 and 100 kDa. The absence of additional signals detected in the precipitated supernatant samples, each corresponding to 1.5 mL of culture supernatant, is likely due to the lower protein amount in the analysed 40 µL of untreated supernatant, which fell below the detection limit.

In biomedical applications, collagen-polyphenol cross-linked hydrogels can improve cell viability, mechanical properties, hydrophilicity, and enzyme resistance while supporting wound healing (Shavandi et al. 2018; Manjari et al. 2020; Han et al. 2024). These results suggest that potential collagen-polyphenol interactions in recombinant collagen polypeptide from moss bioreactors, as well as higher-order aggregates of the collagen polypeptide, could offer additional benefits for cosmetic applications. Further studies are needed to confirm whether such cross-links are formed under our production conditions. Current work is focussed on the bioactivity of the product described here.

Mass spectrometric analysis of the secreted recombinant collagen polypeptide additionally confirmed its prolyl-hydroxylation, a hallmark of collagen biomimicry. The performance of this PTM on recombinant collagens by Physcomitrella is unique compared to other established non-mammalian biotechnological production hosts such as *E. coli* (Kersteen et al. 2004; Pinkas et al. 2011; Shi et al. 2017), yeast (Myllyharju et al. 2000; Myllyharju 2009; Chan et al. 2012), insect cells (Lamberg et al. 1996; Pihlajamaa et al. 1999), and vascular plant systems (Ruggiero et al. 2000; Perret et al. 2001; Ritala et al. 2008; Eskelin et al. 2009; Zhang et al. 2009b). In contrast to these production hosts, which either do not perform prolyl-hydroxylation naturally or achieve only low yields, Physcomitrella, with its six endogenous prolyl-4-hydroxylase (P4H) enzymes (Parsons et al. 2013; Rempfer et al. 2024), was able to facilitate prolyl-hydroxylation of the recombinantly produced collagen polypeptide without requiring additional metabolic engineering, e.g. the expression of heterologous non-plant *P4H* genes.

In the secreted collagen polypeptide fraction, a total of 41 proline residues were detected within the 58% sequence coverage obtained from seven mass spectrometric measurements. Of these identified proline sites, 23 (∼56%) were found to be hydroxylated across the seven datasets, with site-specific hydroxylation prevalences ranging from 10% to 80%. The detected ratio of hydroxylated to non-hydroxylated proline residues in the recombinant collagen polypeptide is consistent with levels found in native human collagens (Guo et al. 2025). Of these 23 hydroxylated prolines, 17 were located at the Yaa position of the Gly-Xaa-Yaa motif, which is typical for native human collagens (Guo et al. 2025). The prolyl-hydroxylation at this position is essential for collagen’s structural integrity, thermal stability, and biological function and also displays the most frequent site of prolyl-hydroxylation in human type III collagen (Shoulders and Raines 2009; Xu et al. 2019; Taga et al. 2021). Among these sites, the integrin-binding motif GAOGER was detected in its hydroxylated form with a prevalence of ∼80%, whereas the second motif, GMOGER, showed no evidence of hydroxyproline modification in the analysed samples. These findings may be linked to differences in the surrounding sequence context, as alanine is the most frequently observed aa preceding hydroxylated prolines in Physcomitrella, while methionine has never been found in this position in previous studies in plant-made biopharmaceutical proteins (Gomord et al. 2010; Rempfer et al. 2024). The remaining six hydroxyprolines were located at the Xaa position of the Gly-Xaa-Yaa motif, a position less frequently hydroxylated in native human collagens (Van Huizen et al. 2019). When hydroxylation at this position occurs, its prevalence typically ranges between 10% and 80%, depending on the specific site, collagen type, and tissue of origin (Kirchner et al. 2021).

Hydroxyprolines in collagens contribute to receptor binding (Chin et al. 2013; Rappu et al. 2019) and, particularly in short peptides derived from collagen, can activate signalling cascades that promote collagen synthesis (Taga et al. 2018), cell proliferation (Ide et al. 2021), hyaluronic acid production (Ohara et al. 2010) and tissue regeneration (Sato et al. 2019). Thus, their presence suggests biomimicry of the collagen polypeptide derived from Physcomitrella and adds potential benefits in terms of bioactivity that are relevant to cosmetic applications. However, further experiments are needed to demonstrate bioactivity *in vitro*.

In photobioreactor cultures, titres of approximately 1 mg/L of secreted collagen polypeptide were achieved, corresponding to about 40-50 g FW (≈ 4-5 g DW) of Physcomitrella protonema. These yields must therefore be interpreted in the context of secretion into the culture medium which differs fundamentally from intracellular accumulation in plant tissues (Doran 2006). First of all, the Physcomitrella culture supernatant contains only minimal amounts of other host proteins (Hoernstein et al. 2018). Furthermore, product secretion in Physcomitrella provides practical advantages over intracellular strategies, as it facilitates scalability and circumvents the laborious extraction and purification steps otherwise required to recover intracellularly stored proteins (Wilken and Nikolov 2012). Beyond these benefits, secretion also ensures passage through the ER and Golgi apparatus, the intracellular sites of key PTMs, such as prolyl-hydroxylation (Parsons et al. 2013; Rempfer et al. 2024).

Direct comparisons of yields across production systems are notoriously difficult. However, based on biomass-specific productivity, our optimized process achieved approximately 32 mg/kg biomass in agitated flask cultures. Compared to other plant-based production systems for recombinant collagens, Physcomitrella clearly outperforms barley cell cultures, where 2-9 µg/kg of product were obtained (Ritala et al. 2008), and even surpasses production in tobacco leaves (0.5-1 mg/kg; Merle et al. 2002) and maize seeds (3-4 mg/kg; Zhang et al. 2009b; Xu et al. 2011). Other plant-based systems, such as maize (Zhang et al. 2009a), barley (Eskelin et al. 2009), and tobacco (Ruggiero et al. 2000), reported intracellular accumulation levels between 13 and 45 mg/kg tissue, which is in the range of our system, while higher yields, up to 200 mg/kg in tobacco leaf tissue, were described by Stein et al. (2009). However, production in these systems requires down-stream processing of the product with potential product losses during purification, besides the cost aspects, whereas the secretion-based production established here does not require extensive down-stream processing. Another advantage of the moss system is its GMP-compliant scalability, with up to 500 L bioreactors in use (Michelfelder et al. 2017).

Prior to further formulation, a three-stage filtration process was developed to effectively separate the collagen polypeptide-containing culture supernatant effectively from plant material and debris. A first filtration through 100 µm sieves allowed the recovery and recycling of Physcomitrella biomass for further processes. A second filtration through 11 µm sieves prevented clogging of subsequently used filters of smaller pore sizes. Among several materials tested for this final clarification, PVDF and PES filters with 5 µm pores achieved nearly 100% recovery, compared to the control filtered through 100 µm and 11 µm mesh sieves. PVDF filters with a pore size of 0.45 µm still recovered 87% of the initial product, whereas PES filters with smaller pore sizes resulted in poor recoveries of 50-60%. In contrast, the other tested filter materials, wwPTFE (0.45 µm), glass microfiber (1.6 µm), cellulose acetate (0.45 µm), nylon (0.45 µm), and polyamide (0.45 µm), resulted in recovery rates below 25%. These results are consistent with the low protein binding properties of PVDF and PES membranes compared to the other materials (Dunleavy 2024). Based on these results, PVDF filters with a pore size of 0.45 µm proved to be the most promising option for integration into the production process, as they achieved the highest product recovery while maintaining a strict filtration limit.

The collagen-rich Physcomitrella culture supernatant is currently being evaluated for efficacy, focusing on the modulation of collagen-related gene expression, ECM remodelling, skin regeneration, and hyaluronic acid release in cell-based assays. If these results are positive, the recombinant human type III collagen polypeptide derived from Physcomitrella can be used as a novel ingredient in various cosmetic formulations.

## Conclusion

This study demonstrates the potential of Physcomitrella as a sustainable production platform for the growing demand for vegan collagen. The recombinant human type III collagen polypeptide produced in Physcomitrella fulfils an important quality criterion through prolyl-hydroxylation, while the established secretion-based, GMP-compliant process in scalable photobioreactors enables the generation of collagen polypeptide-rich supernatants. These supernatants are currently being evaluated for their activity in relation to skin appearance. Overall, these achievements underscore Physcomitrella as a high-performance production host that combines excellent product quality with ethical and sustainability benefits.

## Materials and methods

### Plant material and cell culture

Physcomitrella plants (new species name: *Physcomitrium patens* (Hedw.) Mitt.; IMSC accession number 41269) were cultivated in modified Knop medium (Reski and Abel 1985) according to Decker et al. (2015) and Hoernstein et al. (2018).

### Hydropathicity analysis

To select a soluble human type III collagen polypeptide region suitable for production, the hydropathicity of the amino acid sequence of the human collagen α1(III) chain (P02461) was analysed *in silico* using ExPASy ProtScale (https://web.expasy.org/protscale/). For the analysis the hydropathy scale provided by Kyte and Doolittle (1982) was employed.

### Codon optimization and splice-site avoidance

The sequence encoding for the human type III collagen (COL3A1; NM_000090.4) was obtained from NCBI (https://www.ncbi.nlm.nih.gov/) The codons in the selected collagen polypeptide-encoding sequence were adapted *in silico* towards the Physcomitrella codon usage (Hiss et al. 2017; Top et al. 2021). This codon optimization was performed by Thermo Fisher Scientific. Further, nucleotides in the 11 identified heterosplicing recognition motifs AGGT (Top et al. 2021) were replaced without a change in the respective collagen amino acid sequence (**Supplementary Figure S1**). To possibly enhance production even further, the eight underrepresented codons for Cys, Glu, The, His, Lys, Asn, Gln, and Tyr were replaced with the overrepresented alternative codons (**Supplementary Table S1**) using the online tool physCO (Top et al. 2021; available at www.plant-biotech.uni-freiburg.de). A subsequent search for potential microRNA target sites (Khraiwesh et al. 2010) which was performed with the online tool psRNATarget (https://www.zhaolab.org/psRNATarget/analysis?function=2) yielded no hits. In total, this adaption led to the exchange of 192 of the total 1002 bases (**Supplementary Figure S1**) without changing the amino acid sequence.

### Design of the expression vector

To obtain a secreted product, the desired recombinant collagen polypeptide should be provided with an N-terminal secretory pathway targeting signal peptide and additionally feature a C-terminal His-tag for detection and purification. The codon-optimized and splice site-abolished collagen polypeptide encoding DNA sequence (CDS) was therefore fused *in silico* to the PpAP1 (Pp6c5_10120V6.1) signal-peptide CDS (Schaaf et al. 2004) and the CDS of an 8x His-tag. For cloning purposes, the restriction sites of *Xho*I or *Bgl*II were included at the beginning and the end, respectively, and the sequence obtained was synthesized by GeneArt (Thermo Fisher Scientific, Waltham, USA). The synthesized sequence was subsequently blunt-end ligated in pJet 1.2 (Thermo Fisher Scientific) and the correctness of the sequence was confirmed by sequencing (Eurofins Genomics, Ebersberg, Germany). By using the respective restriction enzymes (Thermo Fisher Scientific), the synthesized sequence as well as a previously used expression vector (Niederau al. 2025) were digested and purified on an agarose gel. The corresponding bands of the expression vector backbone and of the collagen polypeptide-encoding sequence were extracted and subsequently ligated using a T4 ligase (Thermo Fisher Scientific). This resulted in the expression vector containing the strong and endogenous Physcomitrella promoter of the PpActin5 gene (Weise et al. 2006; Niederau et al. 2024), the synthesized target sequence followed by a nos terminator and a hpt cassette for hygromycin selection (Decker et al. 2015) of transformed plants (**Supplementary Figure S2**). For protoplast transfection the expression vector was amplified in *E. coli* DH5a cells, purified using the Plasmid Mega Kit (QIAGEN^®^, Hilden, Germany) according to the manufactureŕs instructions and sterilized via ethanol precipitation according to Sambrook and Russell (2006).

### Generation of transgenic moss lines

Collagen polypeptide-producing moss lines were obtained by stable, polyethylene glycol-mediated transfection of Physcomitrella WT protoplasts (Hohe and Reski 2002; Hohe et al. 2004). Transfection was performed with 50 μg of *Hind*III linearized collagen polypeptide expression vector per reaction according to Decker et al. (2015). Subsequent protoplast regeneration (Schween et al. 2003) followed by the selection of stable transformants on Knop agarose plates containing 50 mg/L hygromycin were performed according to Decker et al. (2015).

### Identification by Western blot

Moss colonies which survived the selection procedure were used to inoculate 500 mL Erlenmeyer flasks containing 180 mL Knop medium with microelements (Knop ME; Egener et al. 2002). They were cultivated for at least 12 weeks prior to screening for collagen production under weekly disruption according to Decker et al. (2015).

To analyse the lines for collagen polypeptide production, protonema suspension cultures were inoculated with 100 mg DW/L and after seven days of cultivation under standard conditions harvested by vacuum filtration. Subsequently, total soluble protein was extracted and purified by His trap spin-down columns (Cytiva, Marlborough, USA) and finally analysed by immunodetections. For this, 100 mg protonema fresh weight (FW) was transferred to 2 mL tubes with one tungsten carbide (Ø 3 mm, QIAGEN^®^, Hilden, Germany) and one glass bead (Ø 3 mm, Roth, Karlsruhe, Germany), frozen in liquid nitrogen and subsequently disrupted by using a tissue lyser (MM 400, Retsch, Haan, Germany) at 30 Hz for 90 seconds. For extraction, 600 µL of extraction buffer (75 mM Na_2_HPO_4_x2H_2_O, 0.5 M NaCl, 20 mM imidazole, 0.05% Tween 20, 10% glycerol, pH 7) were added, the samples were vortexed for 5 min, and subsequently incubated in a cold ultrasound bath for 15 min (Sonorex RK52, Bandelin, Berlin, Germany), followed by 10 min of centrifugation at 14,000 rpm and 4°C. Afterwards, the supernatant was loaded to the His trap spin-down column (Cytiva) which was handled according to the manufacturer’s instructions. Elution of the columns was performed with 200 µL of elution buffer (100 mM Na_2_HPO_4_x2H_2_O, 0.5 M NaCl, 500 mM imidazole, 10% glycerol, pH 7.4). For Western blot analysis, a final concentration (f.c.) of 50 mM dithiothreitol (DTT, Thermo Fisher Scientific) and a f.c. of 1x Laemmli buffer (Bio-Rad, Feldkirchen, Germany) were added and the samples were incubated at 90 °C for 20 min. Subsequently, 10% SDS polyacrylamide gel electrophoresis (SDS-PAGE Ready Gel^®^ Tris-HCl Precast Gels, BioRad) was run at 120 V for 1.5 h and blotted to polyvinylidene fluoride (PVDF) membrane (Amersham^TM^ Hybond P 0.45 µm pore size PVDF blotting membrane, Cytiva) in a Trans-Blot^®^ SD Semi-Dry Transfer Cell (BioRad) for 1h 15 min with 1.5 mA/cm^2^ membrane. The membrane was blocked for 1 h at room temperature in TBS and 0.1% Tween 20 (TBST) containing 4% Amersham^TM^ ECL Prime Blocking Reagent (Cytiva). Subsequently, the membrane was incubated over night with the 1:2,000 in TBST diluted monoclonal anti-6x-His Tag antibody (HIS.H8, MA1-21315, Invitrogen, Thermo Fisher Scientific) at 4 °C. Afterwards, the membrane was washed three times with TBST and incubated with horseradish peroxidase-linked anti-mouse IgG (NA931V, Cytiva) at a dilution of 1:10,000 for 1 h at RT. The blot was washed again and proteins were detected using the Amersham^TM^ ECL Prime Western Blotting Detection Reagent (Cytiva) according to the manufacturer’s instructions. For detection of arabinogalactans, immunodetection with the LM6-M anti-1,5-α-L-arabinan antibody (Plant Probes, Leeds, United Kingdom) was performed as described previously (Bohlender et al. 2022).

### ELISA-mediated collagen quantification

The intracellular accumulation as well as the concentration of collagen polypeptide in the suspension culture supernatant were quantified by ELISA, with identically treated WT analysed in parallel. For intracellular collagen polypeptide analysis, 30 mg of vacuum-filtered protonema material from suspension cultures were homogenized as described for Western blot analysis and extracted in 120 μL of extraction buffer (408 mM NaCl, 60 mM Na_2_HPO_4_, 10.56 mM KH_2_PO_4_, pH 7.4, and 1% protease inhibitor (P9599, Sigma-Aldrich, Merck, Darmstadt, Germany) by vortexing for 5 min, followed by sonication for 15 min in an ultrasonic bath (Bandelin). After centrifugation for 10 min at 14,000 rpm and 4 °C, the supernatants were diluted (1:1,000-1:5,000) in coating buffer (50 mM carbonate/bicarbonate buffer, pH 9.6). A 6xHis-elastin standard protein (ABIN7081465; 28 kDa; antibodies online, Aachen, Germany) was serially diluted in coating buffer (64, 32, 16, 8, 4, 2, 1, and 0 ng/mL). For ELISA, 100 µL of either samples or standards were applied per well of a microtiter plate (NUNC Maxisorb, Wiesbaden, Germany) and incubated overnight at 4 °C. Blocking, incubation, washing, and detection were performed as described previously (Büttner-Mainik et al. 2011), using an anti-6xHis tag monoclonal antibody (HIS.H8, MA1-21315, Invitrogen, Thermo Fisher Scientific; 1:2,000) as primary and an anti-mouse HRP conjugate (NA931V; Cytiva; 1:2,000) as secondary antibody. Under these conditions, the ELISA displayed a linear detection range from 0 to 64 ng/mL. For quantification of collagen polypeptide in the suspension culture supernatant, the ELISA was performed as described above, except that culture supernatants were directly diluted (1:10-1:50) in coating buffer before overnight coating at 4 °C. In both cases, collagen polypeptide concentrations were determined based on the standard curve generated with the serially diluted His-tagged reference protein, with values corrected for background signal and normalized by subtracting the corresponding WT control.

### Secretion optimization

To investigate the accumulation of recombinant collagen polypeptide in the culture supernatant under different cultivation conditions, 200 mL suspension cultures were inoculated with protonema material of line C5 at a density of 300 mg DW/L in aerated 500 mL Erlenmeyer flasks and cultivated under standard conditions, unless otherwise specified. For each condition, WT controls were treated identically and analysed in parallel. Sampling was performed by harvesting 2 mL of suspension culture at the indicated time points. Cell-free supernatants were distributed into 100 µL aliquots, frozen in liquid nitrogen, and stored at -80 °C until ELISA-based analysis.

To test elevated CO_2_ conditions while otherwise maintaining standard cultivation parameters, the corresponding cultures were incubated in a temperature-controlled, illuminated CO_2_ shaker (ISF1-X Climo-Shaker, Kuhner, Basel, Switzerland) set to an atmosphere of 20,000 ppm CO_2_ (2% CO_2_) To investigate the effect of pH, the medium was buffered either with 2.5 mM ammonium tartrate (pH 4.5) or with 1 g/L MES (pH 5.5 and 6.5). Media were adjusted to the respective pH values using 1 M KOH or 1 M HCl and sterile-filtered prior to cultivation. Collagen polypeptide accumulation was quantified by ELISA as previously described. All values were blank-corrected and normalized by subtracting the corresponding WT signals.

### Precipitation of collagen polypeptide

To analyse the quality of the secreted collagen polypeptide, culture supernatant samples were precipitated using acetone. For this, 1-6 mL of culture supernatant were transferred into 50 mL Oak Ridge-style Teflon tubes (Thermo Fisher Scientific) and mixed with five-fold of ice-cold acetone containing 0.2% DTT (Thermo Fisher Scientific). After incubating overnight at -20 °C, the samples were centrifuged at 12,000 x g for 45 min at 0 °C. The supernatant was then discarded, and the pellet was washed with 20 mL of ice-cold acetone, followed by centrifugation under the same conditions. After discarding the supernatant, the pellets were resuspended in 2 mL of acetone, transferred to 2 mL reaction tubes, and centrifuged again under the same conditions. Finally, the supernatant was discarded, and the remaining pellets were air-dried for about 20 minutes. The pellets were stored at -20 °C until further use.

### Mass spectrometric analysis

To investigate the sequence and post-translational modifications of the collagen polypeptide produced, both intracellularly accumulated and secreted fractions were analysed by mass spectrometry. Intracellular collagen polypeptide was prepared as described for the Western blot analysis, whereas secreted collagen polypeptide was obtained by acetone precipitation of the culture supernatant as outlined above. The resulting pellets were dissolved in buffer containing 4% SDS, 125 mM Tris-HCl (pH 6.8), and 50 mM DTT by incubation at 95 °C for 20 min under shaking conditions, followed by centrifugation at 12,000 × g for 5 min at room temperature. For both intracellular and secreted fractions, the protein samples were loaded in duplicate onto two identical 10% SDS-PAGE gels. After electrophoresis, one gel was stained with PageBlue^®^ (Thermo Fisher Scientific), while proteins from the second gel were transferred to a PVDF membrane (Cytiva) and subjected to immunodetection as described above. Bands in the PageBlue^®^-stained gel corresponding to the immunodetected signals were excised and destained as previously outlined in Bohlender et al. (2020). Overnight digestion was carried out with 0.2 µg of trypsin (Trypsin Gold, Promega, Wisconsin, USA) at 37 °C. The resulting peptides were recovered and desalted using C18 StageTips (Thermo Fisher Scientific), followed by analysis on Orbitrap Elite and Orbitrap XL mass spectrometers (Thermo Fisher Scientific), as previously described (Njenga et al. 2024).

The raw data of the measurement of intracellular collagen were processed using Mascot Distiller V2.7.10, and a database search was conducted with the Mascot server V2.7 (Matrix Science, London, United Kingdom). Processed spectra were searched against a database containing all Physcomitrella protein models (V3.372; Lang et al. 2018), the sequence of the collagen polypeptide, and a database of known contaminants (269 entries; Mueller et al. 2014), using a precursor mass tolerance of 8 ppm and a fragment mass tolerance of 0.6 Da. Variable modifications included Gln→pyroGlu (N-term. Q) -17.026549 Da; dehydration Glu→pyroGlu (N-term. E) - 18.010565 Da; oxidation +15.994915 Da (P and M); deamidation +0.984016 Da (N); the attachment of one to five arabinoses to hydroxylated prolines +148.037173 Da (Ara), +280.079432 Da (Ara_2_), +412.121691 Da (Ara_3_), +544.163950 Da (Ara_4_), +676.206209 Da (Ara_5_), (Hyp_Ara_1-5_) and the attachment of one hexose to hydroxylated prolines +178.047738 Da (Hyp_Hex).

For label-free quantitation of identified Hyp sites, the seven measurements from secreted samples were run again in *FragPipe* v23.1 configured for data dependent acquisition (DDA). Peptides were identified using *MSFragger* (Kong et al. 2017) and a Physcomitrella protein database (48,582 entries; Lang et al. 2018) including target and decoy sequences for FDR estimation as well as the sequence of the collagen polypeptide. A precursor mass tolerance of -20 to +20 ppm was used, enzyme specificity was set to semi trypsin and a peptide mass range of 500 to 5,000 Da with a minimum peptide length of 7 and a maximum of 40 amino acids was set. Carbamidomethylation of cysteines (+57.02146 Da) was set as fixed modification. Variable modifications, oxidation (+15.9949 Da) of methionine (M) and proline (P), cyclisation of N-terminal asparagine and carbamidomethylated cysteine (-17.0265 Da) as well as deamidation of glutamine (Q, -18.010565 Da) and acetylation at the protein N-terminus (+42.010567 Da). Validation of peptide-spectrum matches, and the prolyl-hydroxylation site was done with *MSBooster* (Yang et al. 2023), Percolator (Käll et al. 2007; The et al. 2017), and PTMProphet (Shteynberg et al. 2019). Quantification was conducted using *IonQuant* applying a protein-level false discovery rate (FDR) threshold of 0.01.

### Dry weight measurement

To determine the dry weight (DW) of suspension cultures from flasks, three samples of 10 mL each were collected under shaking conditions, filtered through pre-dried and pre-weighed Miracloth tissues (Merck, Darmstadt, Germany), and subsequently dried for 2 h at 105 °C. The DW was then determined using an analytical balance (CPA 3245; Sartorius, Göttingen, Germany). For bioreactor cultures, 100 mL of suspension were transferred into a beaker equipped with a magnetic stirrer. While stirring, triplicate samples of 10–20 mL were collected and processed as described above to determine the culture dry weight. (DW).

### Bioreactor operation

For large-scale recombinant collagen polypeptide production, 5 L stirred-tank photobioreactors (Applikon Biotechnology, Schiedam, The Netherlands) were inoculated with 400 mg DW of moss protonema material per litre medium. Aeration was maintained at 1.5 vvm using CO_2_-enriched air (2% CO_2_, 0.3 vvm), and agitation was provided by a three-blade pitched impeller operating at 375 rpm. The pH was controlled at 4.8, and the temperature was kept constant at 22 °C. Continuous illumination was supplied by twelve neutral white (4000 K) LED strips (MaxLine70, Lumitronix, Hechingen, Germany), evenly distributed across the bioreactor surface. Light intensity (µmol m^-2^ s^-1^) was measured and calibrated with a light meter (LI-250A, LI-COR, Bad Homburg, Germany) at the inner bioreactor wall prior to sterilization. The applied light regime was as follows: light intensity was initially set to 100 µmol m^-2^ s^-1^ and increased to 350 µmol m^-2^ s^-1^ once a biomass concentration of 2 g DW/L was reached. When biomass reached 2.8 g DW/L, the intensity was further increased to 680 µmol m^-2^ s^-1^ and maintained at this level until harvest, which was performed at an accumulated biomass of 4 g DW/L.

### Bioreactor sampling

During the bioreactor runs, samples were collected to monitor biomass accumulation and to determine both intracellular and secreted collagen polypeptide concentrations. For sampling, 100 mL of suspension culture was harvested from the bioreactor and replaced with fresh medium. The collected suspension was stirred in a beaker and first 3x 10 mL samples for dry weight determination were collected. For ELISA measurements, 100 µL aliquots were collected after allowing the suspension to sediment for 10 min, in order to minimize the carryover of moss tissue. Aliquots were frozen immediately in liquid nitrogen and stored at -80 °C until analysis. For protonema sampling, the remaining suspension culture was vacuum filtered, and 30 mg as well as 100 mg fresh weight (FW) of protonema was transferred into 2 mL reaction tubes, frozen in liquid nitrogen, and stored at −80 °C until further use.

### Filtration of collagen-containing supernatants

To remove residual cells and debris from collagen polypeptide-containing culture supernatants, which had already been separated from moss tissue using 100 µm and 11 µm protoplast sieves, the suitability of various filter materials was evaluated. The tested filters included cellulose acetate (0.45 µm; Ø 50 mm; OE67; Whatman^TM^; Cytiva), glass microfiber (1.6 µm; Ø 50 mm; GF/A; Whatman^TM^; Cytiva), nylon (0.45 µm; Ø 47 mm; Nylaflo^TM^; Cytiva), polyethersulfone (PES; 0.22 µm; Ø 33 mm; Rotilabo^®^, Roth; Karlsruhe, Germany and 1 µm; Ø 25mm; Whatman^TM^ Puradisc; Cytiva and 5 µm; Ø 25mm; Chromafil^®^Xtra; Macherey-Nagel, Düren, Germany), polyamide (0.45 µm; Ø 50 mm; NL17; Whatman^TM^; Cytiva), polyvinylidene difluoride (PVDF; 0.45 µm; Ø 33 mm, Rotilabo^®^, Roth; and 5 µm; Ø 47 mm; Durapore^®^; Merck), and water-wettable polytetrafluoroethylene (wwPTFE; 0.45 µm; Ø 47 mm; Cytiva). These filters were used either as disc filters, mounted in an individual filter holder (16201, Sartorius) and operated with a vacuum flask, or, in the case of PES (0.22 µm, 1 µm, and 5 µm) and PVDF (0.45 µm), as syringe filters. For each case, 100 mL of harvested bioreactor supernatant were processed. Collagen polypeptide concentrations in the filtrates and in the unfiltered control were quantified by ELISA, and recoveries were calculated as percentages of the initial collagen polypeptide concentration present prior to filtration.

### Nucleic acids extraction and quantitative PCR

Both genomic DNA (for qPCR) and RNA (for qRT-PCR) were extracted from 100 mg of protonema material frozen in liquid nitrogen (Schlink and Reski 2002). Protonema material for qRT-PCR analyses was harvested from equally inoculated flasks after seven days of cultivation. For the qPCR all subsequent steps were performed as described in Ruiz-Molina et al. (2022b), while the qRT-PCR analyses were conducted as carried out in Bohlender et al. (2020). Primers were designed using the online tool Primer3Plus (https://www.primer3plus.com/) with the qPCR server settings, and checked for specificity against the Physcomitrella genome in the Phytozome v13 database (https://phytozome-next.jgi.doe.gov) (Goodstein et al. 2012). Primer specificity and efficiency were validated for each primer pair using DNA serial dilutions. For construct copy number determination, primer pairs targeting the collagen polypeptide CDS, the Actin5 promoter, and the hpt selection cassette were used. Reference signals for single integrations for normalization were obtained with two primer pairs (CLF5 and CLF7) targeting the single-copy gene PpCLF (Pp6c22_9280V6.1, Pereman et al. 2016). Transgene copy numbers were determined using two control calibrators: the Bag4.2 moss line, which carries a single integration of the hpt selection cassette, and the WT, which served as a single-copy reference for the Actin5 promoter. For qRT-PCR, a primer pair targeting the collagen polypeptide CDS was used, and normalization was performed with two primer pairs amplifying regions of the strongly expressed housekeeping genes LWD (Pp6c22_10850V6.1, Yin et al. 2021) encoding the WD repeat-containing protein 68 and L21 (Pp6c13_1120V6.1; Beike et al. 2015) encoding the large subunit ribosomal protein L21. All primers were synthesized by Eurofins Genomics and are listed in **Supplementary Table S2**.

## Acknowledgements

We thank Christine Glockner and Britta Rothgänger for technical assistance and Anne Katrin Prowse for proof reading.

## Author contributions

L.L.B., A.M. and J.P. performed research. L.L.B., J.P., S.N.W.H., G.G., B.H., E.L.D. and R.R. analysed data. E.L.D. and R.R. supervised research. B.H., E.L.D. and R.R. acquired funding. L.L.B. and R.R. wrote the manuscript. All authors read and approved the final version of the manuscript.

## Data availability

The Physcomitrella WT used in this study, as well as the collagen-producing transgenic strains are available from the International Moss Stock Center IMSC (https://www.moss-stock-center.org) under the accession numbers 41269 (WT D14), 40996 (C5), and 40997 (C6.1). The mass spectrometry proteomics data have been deposited at the ProteomeXchange Consortium via the PRIDE partner repository (Deutsch et al. 2023; Perez-Riverol et al. 2025) with the dataset identifiers PXD071319 and PXD071359.

## Supplementary Information

**Supplementary Figure S1.**
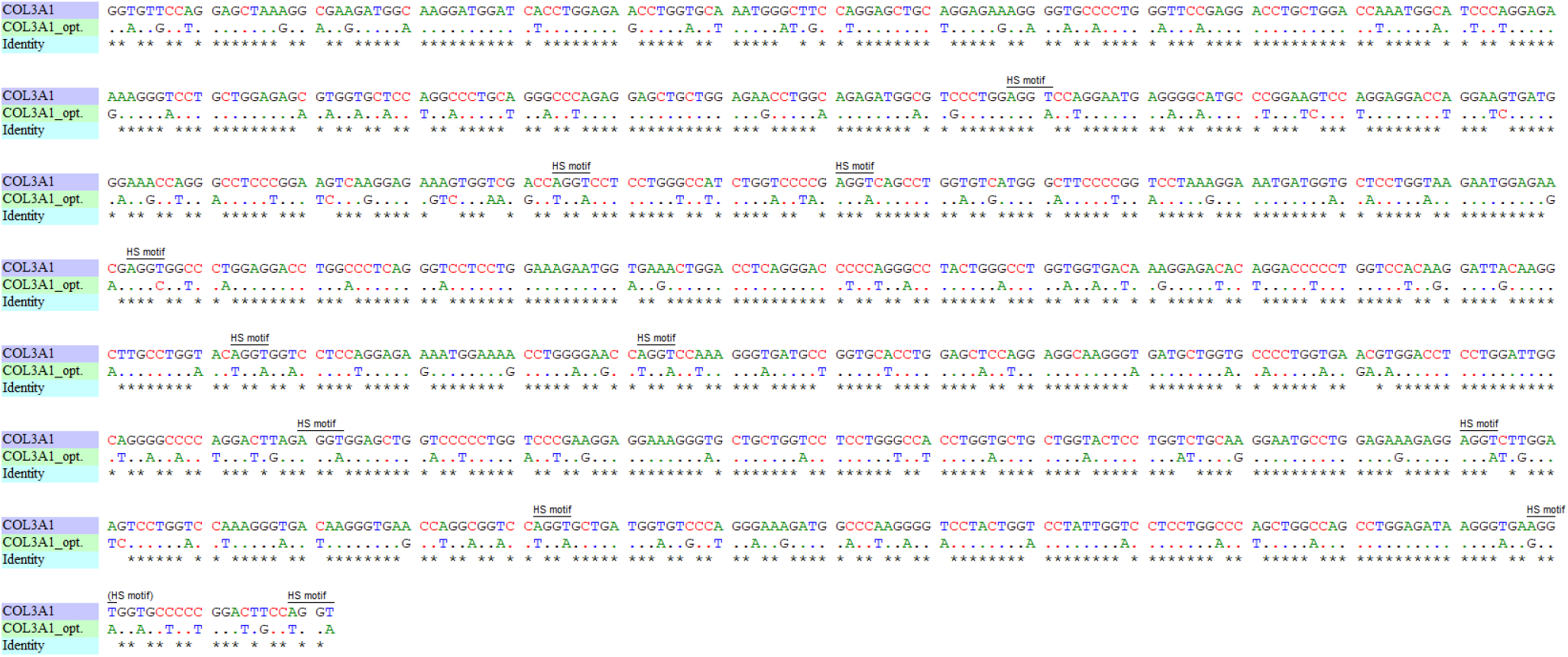
Alignment of the selected coding region (nucleotides 1266–2367) of the human COL3A1 CDS (COL3A1, NM_000090.4) and the codon- and splice site-optimized COL3A1 sequence (COL3A1_opt.) used for generation of collagen-producing Physcomitrella lines. The CDS of the human collagen α1(III) chain (COL3A1) was obtained from NCBI (NM_000090.4). The region coding for the selected recombinant collagen polypeptide is depicted, aligned with the corresponding sequence after *in silico* codon and potential heterosplice sites optimization. Codon optimization according to Physcomitrella codon usage was carried out by Thermo Fisher Scientific. In addition, underrepresented codons (as listed in **Supplementary Table S1**) were substituted with corresponding, overrepresented alternatives using the physCO online tool (available at www.plant-biotech.uni-freiburg.de). To prevent potential heterosplicing, nucleotides within the heterosplicing recognition motif AGGT (HS motif, Top et al. 2021) were silently mutated *in silico*. The resulting optimized sequence (COL3A1_opt.) encodes the original amino acid sequence and was modified exclusively at the nucleotide level.

**Supplementary Figure S2.**
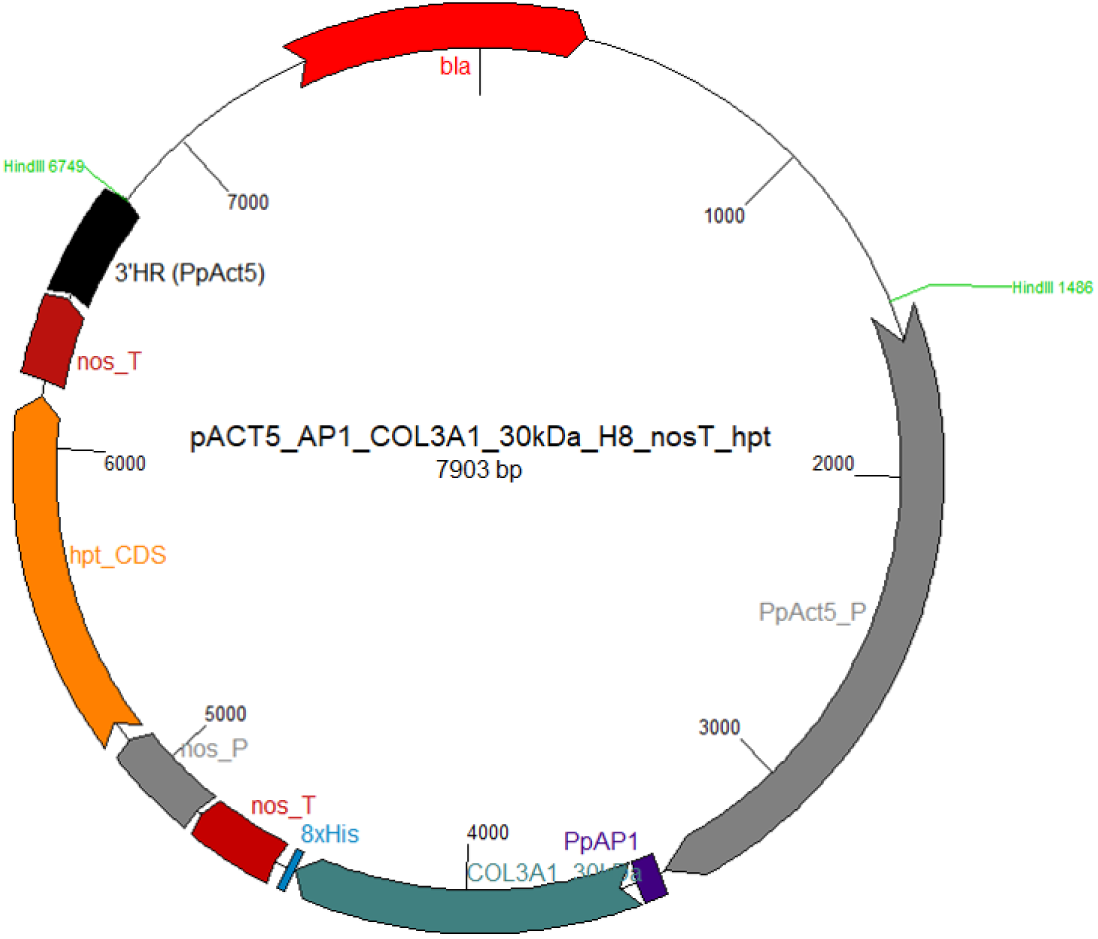
Schematic representation of the pJET1.2-based vector containing the expression construct used to generate recombinant collagen-polypeptide producing Physcomitrella lines. The codon- and splice-site optimized collagen polypeptide CDS including the CDS for the aspartic protease signal peptide PpAP1 (Pp6c5_10120V6.1; Schaaf et al. 2004) and an 8xHis-tag were cloned into an pJET1.2-based expression vector containing the Physcomitrella Actin5 (PpActin5) promoter (Weise et al. 2006; Niederau et al. 2024) and the nos terminator (nos_T). For selection purposes this vector additionally contained a hygromycin selection cassette coding for a hygromycin B phosphotransferase (hpt; Decker et al. 2015) under the control of a nos promoter (nos_P) and a nos terminator. Targeted genome integration via homologous recombination was facilitated by a 327 bp long 3’ homologous region of the 3’ UTR of the endogenous actin 5 encoding gene, while the Actin5 promoter sequence serves as 5’ homologous flank. Prior to transformation the plasmid was linearized using *Hin*dIII.

**Supplementary Figure S3.**
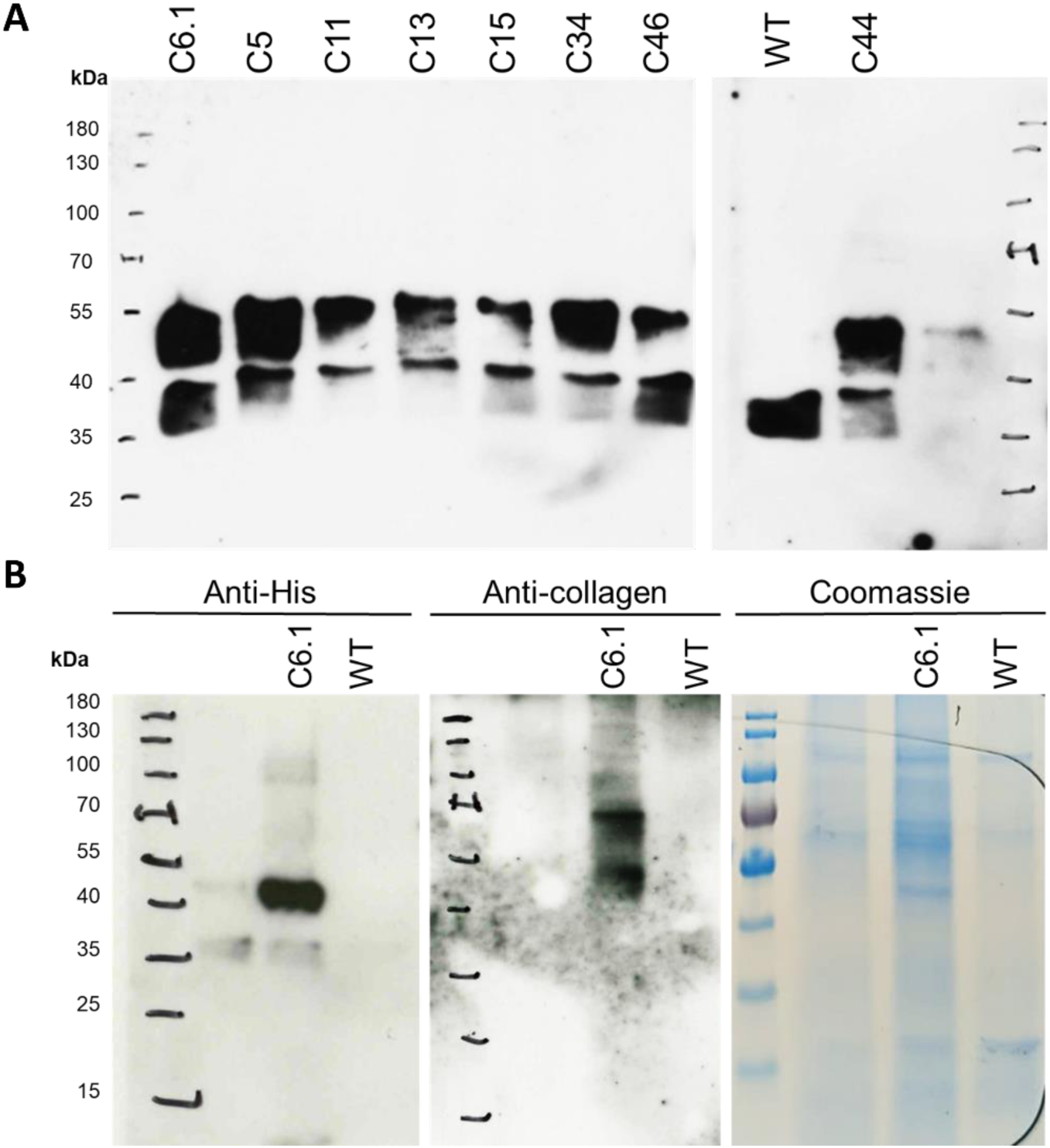
Uncropped immunodetection images and Coomassie stained SDS-PAGE gel image corresponding to Figure 2. **A** Corresponds to **Figure 2A** and **B** corresponds to **Figure 2B**.

**Supplementary Figure S4.**
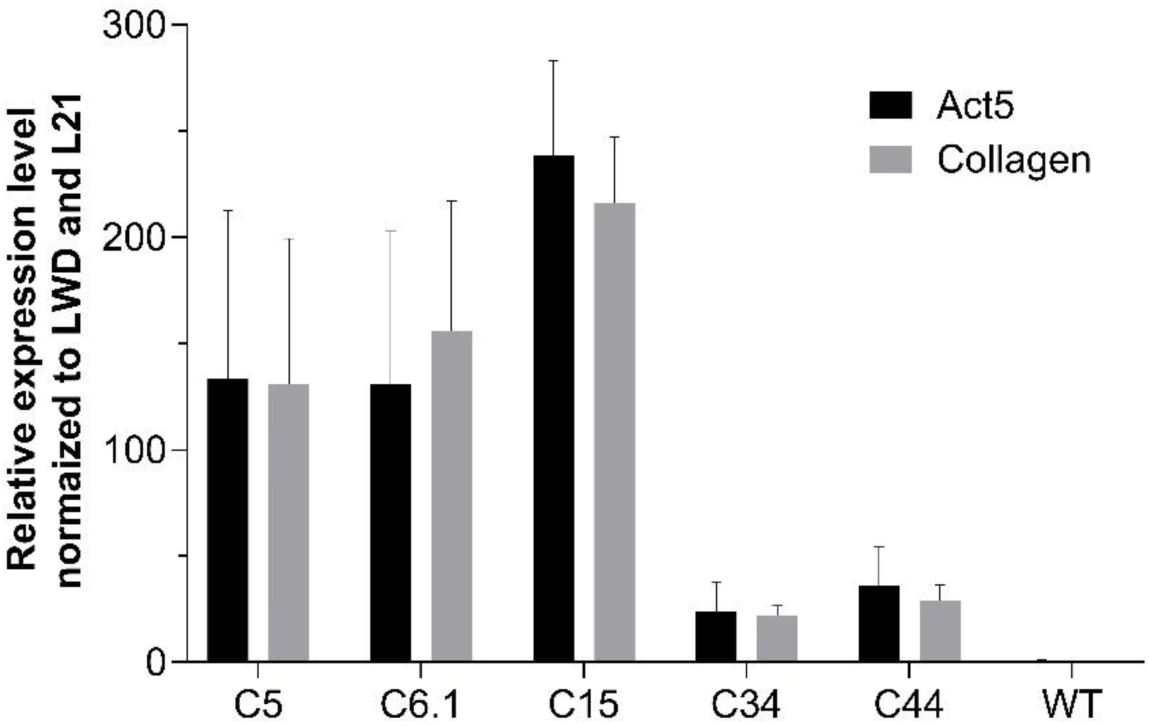
Quantitative analysis of construct copy numbers in selected moss lines. The five selected collagen polypeptide-producing lines and an equally treated WT control were investigated for the number of integrated expression constructs by qPCR. Copy numbers were analysed using the primer pairs targeting the Actin5 promotor (Act5) and the collagen CDS (Collagen). Internal normalization to single integration signals was performed with signals obtained by two primer pairs targeting the endogenous single copy CLF gene and single integration signals for the Actin5 promoter were gained from the wild type (WT). Bars represent normalized mean values ± SD from n = 3 technical replicates.

**Supplementary Figure S5.**
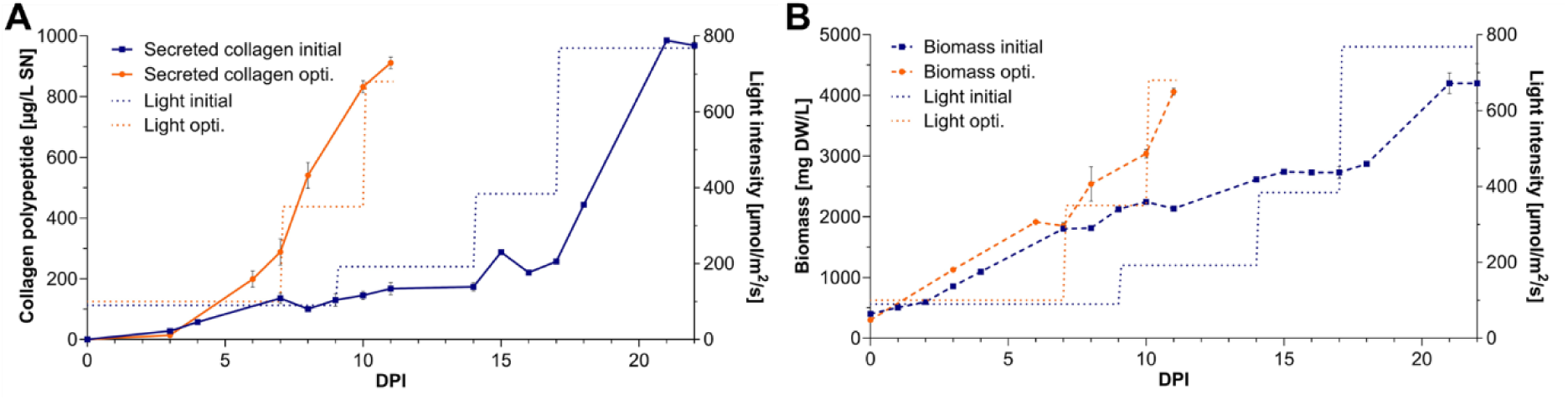
Split version of Figure 5A, depicting parameters during 5 L photobioreactor runs of line C5 under initial and optimized conditions. **A** Secreted collagen polypeptide accumulation profiles (solid lines) and applied light intensity settings (dotted lines). Collagen concentrations were quantified by ELISA (mean ± SD, technical triplicates). **B** Moss biomass accumulation (dashed lines) and applied light intensity settings (dotted lines). Biomass quantification was performed by dry weight measurements (mean ± SD, technical duplicates).

**Supplementary Figure S6.**
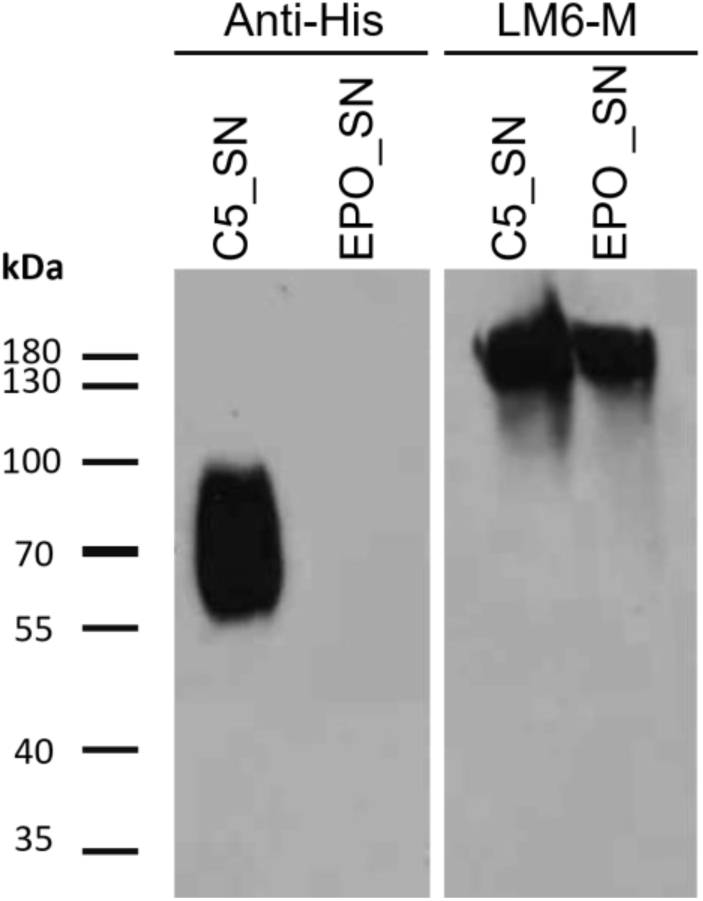
Immunodetection of precipitated bioreactor culture supernatants. Precipitated and blotted bioreactor culture supernatants (SN) from the rhEPO-producing moss line *Δ*galt1 (EPO; Bohlender et al. 2022) and from line C5 were analysed with an anti-His antibody and the anti-1,5-α-L-arabinan antibody LM6-M. While LM6-M detection revealed the presence of arabinogalactan proteins in both samples at an apparent molecular weight of approximately 180 kDa (Lee et al. 2005; Bohlender et al. 2022), no corresponding signals were observed in the 55–90 kDa molecular weight range, where the secreted collagen peptide, as detected by the anti-His antibody, accumulates.

**Supplementary Figure S7.**
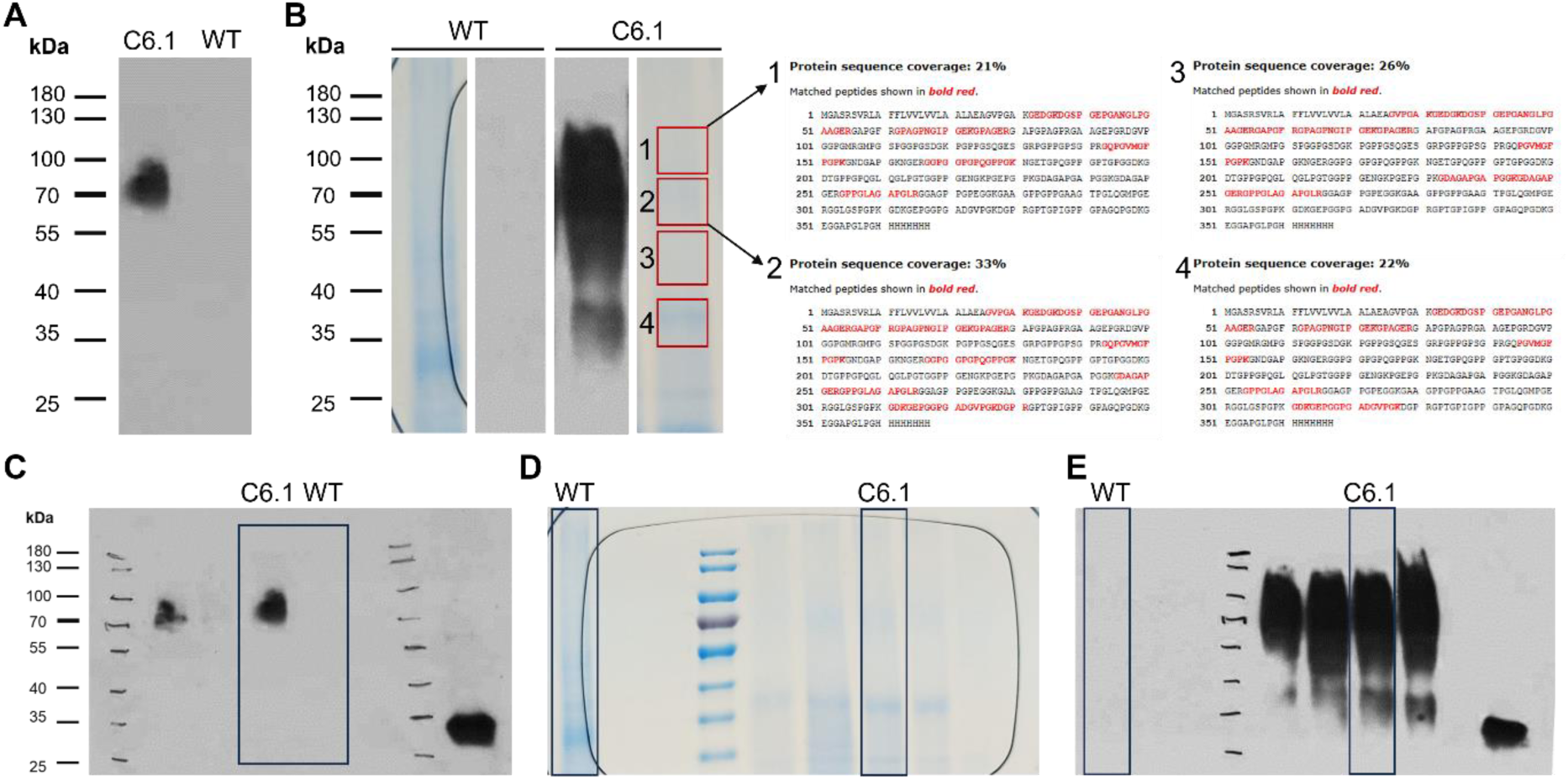
Immunodetection from bioreactor culture supernatants, corresponding SDS-PAGE gels, mass spectrometrically determined sequence coverages of the recombinant collagen polypeptide, and uncropped images. **A** Anti-His antibody-based immunodetection of SDS-PAGE–separated and PVDF-transferred, non-precipitated bioreactor culture supernatants (40 µL per lane) from line C6.1 and the wild type (WT). **B** Coomassie-stained SDS-PAGE gels and corresponding anti-His immunodetection of 1.5 mL acetone-precipitated bioreactor culture supernatants from line C6.1 after 16 days of cultivation, compared to an equally treated WT control. Gel regions corresponding to immunodetection signals (marked as 1-4) were excised and analysed by mass spectrometry. The identified sequence coverages of the recombinant collagen polypeptide are marked in red. **C** Uncropped immunodetection image corresponding to **Supplementary Figure S7A**. **D** Uncropped SDS-PAGE gel image corresponding to **Supplementary Figure S7B**. **E** Uncropped immunodetection image corresponding to **Figure 6B** and **Supplementary Figure S7B**.

**Supplementary Table S1.** Underrepresented codons that were exchanged by overrepresented ones in the optimized CDS used for collagen production. These selections are based the codon usage tables described by Hiss et al. (2017) and Nakamura et al. (2000).

| Amino acid | Underrepresented codon | Overrepresented codon |
| --- | --- | --- |
| Cys | TGT | TGC |
| Glu | GAA | GAG |
| Phe | TTT | TTC |
| His | CAT | CAC |
| Lys | AAA | AAG |
| Asn | AAT | AAC |
| Gln | CAA | CAG |
| Tyr | TAT | TAC |

**Supplementary Table S2.**
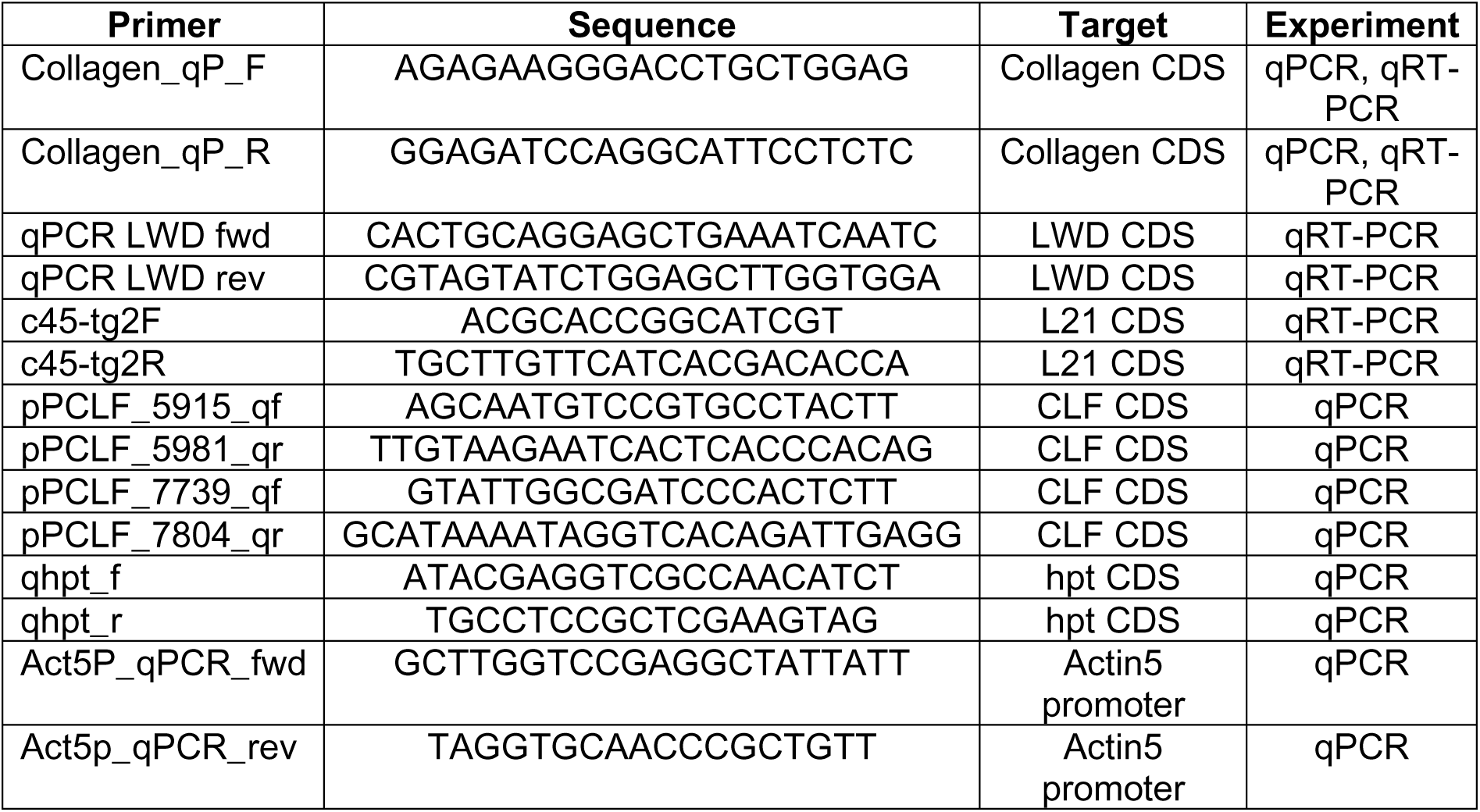
Primers used for quantitative PCR.

**Supplementary Table S3.** Quantitation summary of identified hydroxylated prolines in secreted collagen samples. Shown are the proline residues identified across the seven analysed measurements (raw files), including their respective G-X-Y motif and their corresponding amino acid positions in the recombinant collagen polypeptide (Position_Motif). Log₂-transformed peptide intensities were used as quantitative values. These were first normalized to the median intensity within each measurement and subsequently centred to the global median intensity across all measurements. For each identified hydroxyproline or its unmodified counterpart the quantitative values from each corresponding peptide isoform were summed within each measurement (Raw File). The relative proportion of modified versus unmodified prolines per peptide is given as a percentage.

| Raw File | Position Motif | Type | Summed Intensity | Total Intensity | Percent |
| --- | --- | --- | --- | --- | --- |
| ELITE-RSLC037807 | 40_GSP | Mod. | 20.8633 | 84.8853 | 24.58 |
| ELITE-RSLC037807 | 40_GSP | Unmod. | 64.0221 | 84.8853 | 75.42 |
| ELITE-RSLC037807 | 43_GEP | Unmod. | 84.8853 | 84.8853 | 100.00 |
| ELITE-RSLC037807 | 49_GLP | Unmod. | 84.8853 | 84.8853 | 100.00 |
| ELITE-RSLC037807 | 63_GPA | Mod. | 16.1702 | 16.1702 | 100.00 |
| ELITE-RSLC037807 | 66_GPN | Mod. | 16.1702 | 16.1702 | 100.00 |
| ELITE-RSLC037807 | 70_GIP | Unmod. | 34.5415 | 34.5415 | 100.00 |
| ELITE-RSLC037807 | 75_GPA | Unmod. | 35.5944 | 35.5944 | 100.00 |
| ELITE-RSLC037807 | 94_GEP | Unmod. | 18.7694 | 18.7694 | 100.00 |
| ELITE-RSLC037807 | 100_GVP | Mod. | 18.7694 | 53.3520 | 35.18 |
| ELITE-RSLC037807 | 100_GVP | Unmod. | 34.5826 | 53.3520 | 64.82 |
| ELITE-RSLC037807 | 103_GGP | Mod. | 18.7694 | 36.9129 | 50.85 |
| ELITE-RSLC037807 | 103_GGP | Unmod. | 18.1435 | 36.9129 | 49.15 |
| ELITE-RSLC037807 | 145_GQP | Unmod. | 23.2473 | 23.2473 | 100.00 |
| ELITE-RSLC037807 | 153_GPK | Unmod. | 23.2473 | 23.2473 | 100.00 |
| ELITE-RSLC037807 | 250_GAP | Mod. | 17.6325 | 17.6325 | 100.00 |
| ELITE-RSLC037807 | 255_GPP | Unmod. | 20.5607 | 20.5607 | 100.00 |
| ELITE-RSLC037807 | 256_GPP | Unmod. | 61.5526 | 61.5526 | 100.00 |
| ELITE-RSLC037807 | 262_GAP | Mod. | 38.1931 | 38.1931 | 100.00 |
| ELITE-RSLC037807 | 316_GEP | Unmod. | 33.5776 | 33.5776 | 100.00 |
| ELITE-RSLC037807 | 319_GGP | Unmod. | 33.5776 | 33.5776 | 100.00 |
| ELITE-RSLC037807 | 325_GVP | Unmod. | 33.5776 | 33.5776 | 100.00 |
| ELITE-RSLC037809 | 28_GVP | Mod. | 66.4161 | 139.2935 | 47.68 |
| ELITE-RSLC037809 | 28_GVP | Unmod. | 72.8774 | 139.2935 | 52.32 |
| ELITE-RSLC037809 | 40_GSP | Mod. | 133.3001 | 529.5811 | 25.17 |
| ELITE-RSLC037809 | 40_GSP | Unmod. | 396.2810 | 529.5811 | 74.83 |
| ELITE-RSLC037809 | 43_GEP | Unmod. | 599.2329 | 599.2329 | 100.00 |
| ELITE-RSLC037809 | 49_GLP | Mod. | 58.1106 | 599.2329 | 9.70 |
| ELITE-RSLC037809 | 49_GLP | Unmod. | 541.1223 | 599.2329 | 90.30 |
| ELITE-RSLC037809 | 58_GAP | Unmod. | 20.2887 | 20.2887 | 100.00 |
| ELITE-RSLC037809 | 63_GPA | Mod. | 39.3541 | 58.2412 | 67.57 |
| ELITE-RSLC037809 | 63_GPA | Unmod. | 18.8871 | 58.2412 | 32.43 |
| ELITE-RSLC037809 | 66_GPN | Mod. | 39.3541 | 98.2693 | 40.05 |
| ELITE-RSLC037809 | 66_GPN | Unmod. | 58.9152 | 98.2693 | 59.95 |
| ELITE-RSLC037809 | 70_GIP | Unmod. | 176.4821 | 176.4821 | 100.00 |
| ELITE-RSLC037809 | 75_GPA | Unmod. | 176.2793 | 176.2793 | 100.00 |
| ELITE-RSLC037809 | 94_GEP | Mod. | 23.9800 | 66.7997 | 35.90 |
| ELITE-RSLC037809 | 94_GEP | Unmod. | 42.8198 | 66.7997 | 64.10 |
| ELITE-RSLC037809 | 100_GVP | Mod. | 23.9800 | 61.5583 | 38.95 |
| ELITE-RSLC037809 | 100_GVP | Unmod. | 37.5784 | 61.5583 | 61.05 |
| ELITE-RSLC037809 | 103_GGP | Mod. | 23.9800 | 77.9768 | 30.75 |
| ELITE-RSLC037809 | 103_GGP | Unmod. | 53.9969 | 77.9768 | 69.25 |
| ELITE-RSLC037809 | 145_GQP | Mod. | 44.9735 | 93.5253 | 48.09 |
| ELITE-RSLC037809 | 145_GQP | Unmod. | 48.5518 | 93.5253 | 51.91 |
| ELITE-RSLC037809 | 151_GFP | Mod. | 21.0235 | 88.6469 | 23.72 |
| ELITE-RSLC037809 | 151_GFP | Unmod. | 67.6234 | 88.6469 | 76.28 |
| ELITE-RSLC037809 | 153_GPK | Mod. | 21.0235 | 161.6960 | 13.00 |
| ELITE-RSLC037809 | 153_GPK | Unmod. | 140.6725 | 161.6960 | 87.00 |
| ELITE-RSLC037809 | 169_GPG | Unmod. | 17.9621 | 17.9621 | 100.00 |
| ELITE-RSLC037809 | 172_GPG | Unmod. | 17.9621 | 17.9621 | 100.00 |
| ELITE-RSLC037809 | 174_GPQ | Unmod. | 17.9621 | 17.9621 | 100.00 |
| ELITE-RSLC037809 | 177_GPP | Unmod. | 17.9621 | 17.9621 | 100.00 |
| ELITE-RSLC037809 | 178_GPP | Unmod. | 17.9621 | 17.9621 | 100.00 |
| ELITE-RSLC037809 | 241_GAP | Unmod. | 16.0661 | 16.0661 | 100.00 |
| ELITE-RSLC037809 | 250_GAP | Mod. | 68.8330 | 84.9600 | 81.02 |
| ELITE-RSLC037809 | 250_GAP | Unmod. | 16.1270 | 84.9600 | 18.98 |
| ELITE-RSLC037809 | 255_GPP | Unmod. | 90.1647 | 90.1647 | 100.00 |
| ELITE-RSLC037809 | 256_GPP | Mod. | 26.2192 | 160.7007 | 16.32 |
| ELITE-RSLC037809 | 256_GPP | Unmod. | 134.4815 | 160.7007 | 83.68 |
| ELITE-RSLC037809 | 262_GAP | Mod. | 93.2291 | 113.0045 | 82.50 |
| ELITE-RSLC037809 | 262_GAP | Unmod. | 19.7754 | 113.0045 | 17.50 |
| ELITE-RSLC037809 | 283_GPP | Unmod. | 20.1412 | 20.1412 | 100.00 |
| ELITE-RSLC037809 | 285_GPP | Unmod. | 20.1412 | 20.1412 | 100.00 |
| ELITE-RSLC037809 | 286_GPP | Unmod. | 20.1412 | 20.1412 | 100.00 |
| ELITE-RSLC037809 | 292_GTP | Unmod. | 20.1412 | 20.1412 | 100.00 |
| ELITE-RSLC037809 | 298_GMP | Unmod. | 20.1412 | 20.1412 | 100.00 |
| ELITE-RSLC037809 | 307_GSP | Unmod. | 19.6213 | 19.6213 | 100.00 |
| ELITE-RSLC037809 | 309_GPK | Unmod. | 19.6213 | 19.6213 | 100.00 |
| ELITE-RSLC037809 | 316_GEP | Unmod. | 289.2604 | 289.2604 | 100.00 |
| ELITE-RSLC037809 | 319_GGP | Mod. | 21.3337 | 290.5687 | 7.34 |
| ELITE-RSLC037809 | 319_GGP | Unmod. | 269.2350 | 290.5687 | 92.66 |
| ELITE-RSLC037809 | 325_GVP | Mod. | 57.9244 | 309.5610 | 18.71 |
| ELITE-RSLC037809 | 325_GVP | Unmod. | 251.6366 | 309.5610 | 81.29 |
| ELITE-RSLC037809 | 330_GPR | Unmod. | 193.2421 | 193.2421 | 100.00 |
| ELITE-RSLC037842 | 28_GVP | Mod. | 16.5849 | 101.8921 | 16.28 |
| ELITE-RSLC037842 | 28_GVP | Unmod. | 85.3072 | 101.8921 | 83.72 |
| ELITE-RSLC037842 | 40_GSP | Mod. | 39.8071 | 204.3013 | 19.48 |
| ELITE-RSLC037842 | 40_GSP | Unmod. | 164.4942 | 204.3013 | 80.52 |
| ELITE-RSLC037842 | 43_GEP | Mod. | 19.7206 | 266.1995 | 7.41 |
| ELITE-RSLC037842 | 43_GEP | Unmod. | 246.4790 | 266.1995 | 92.59 |
| ELITE-RSLC037842 | 49_GLP | Unmod. | 284.1138 | 284.1138 | 100.00 |
| ELITE-RSLC037842 | 63_GPA | Mod. | 35.2337 | 35.2337 | 100.00 |
| ELITE-RSLC037842 | 66_GPN | Mod. | 35.2337 | 52.2873 | 67.38 |
| ELITE-RSLC037842 | 66_GPN | Unmod. | 17.0536 | 52.2873 | 32.62 |
| ELITE-RSLC037842 | 70_GIP | Unmod. | 103.9241 | 103.9241 | 100.00 |
| ELITE-RSLC037842 | 75_GPA | Mod. | 17.7395 | 87.3611 | 20.31 |
| ELITE-RSLC037842 | 75_GPA | Unmod. | 69.6216 | 87.3611 | 79.69 |
| ELITE-RSLC037842 | 103_GGP | Unmod. | 19.7747 | 19.7747 | 100.00 |
| ELITE-RSLC037842 | 145_GQP | Unmod. | 17.4944 | 17.4944 | 100.00 |
| ELITE-RSLC037842 | 151_GFP | Mod. | 17.4944 | 81.1963 | 21.55 |
| ELITE-RSLC037842 | 151_GFP | Unmod. | 63.7019 | 81.1963 | 78.45 |
| ELITE-RSLC037842 | 153_GPK | Mod. | 17.4944 | 100.1879 | 17.46 |
| ELITE-RSLC037842 | 153_GPK | Unmod. | 82.6935 | 100.1879 | 82.54 |
| ELITE-RSLC037842 | 169_GPG | Unmod. | 31.8294 | 31.8294 | 100.00 |
| ELITE-RSLC037842 | 172_GPG | Unmod. | 31.8294 | 31.8294 | 100.00 |
| ELITE-RSLC037842 | 174_GPQ | Unmod. | 31.8294 | 31.8294 | 100.00 |
| ELITE-RSLC037842 | 177_GPP | Unmod. | 31.8294 | 31.8294 | 100.00 |
| ELITE-RSLC037842 | 178_GPP | Unmod. | 31.8294 | 31.8294 | 100.00 |
| ELITE-RSLC037842 | 250_GAP | Mod. | 21.4815 | 21.4815 | 100.00 |
| ELITE-RSLC037842 | 255_GPP | Unmod. | 66.0051 | 66.0051 | 100.00 |
| ELITE-RSLC037842 | 256_GPP | Mod. | 24.9956 | 87.4866 | 28.57 |
| ELITE-RSLC037842 | 256_GPP | Unmod. | 62.4910 | 87.4866 | 71.43 |
| ELITE-RSLC037842 | 262_GAP | Mod. | 68.8090 | 87.4866 | 78.65 |
| ELITE-RSLC037842 | 262_GAP | Unmod. | 18.6776 | 87.4866 | 21.35 |
| ELITE-RSLC037842 | 282_GPP | Unmod. | 17.7921 | 17.7921 | 100.00 |
| ELITE-RSLC037842 | 283_GPP | Unmod. | 17.7921 | 17.7921 | 100.00 |
| ELITE-RSLC037842 | 285_GPP | Unmod. | 17.7921 | 17.7921 | 100.00 |
| ELITE-RSLC037842 | 286_GPP | Unmod. | 17.7921 | 17.7921 | 100.00 |
| ELITE-RSLC037842 | 292_GTP | Unmod. | 17.7921 | 17.7921 | 100.00 |
| ELITE-RSLC037842 | 298_GMP | Unmod. | 17.7921 | 17.7921 | 100.00 |
| ELITE-RSLC037842 | 316_GEP | Unmod. | 81.4029 | 81.4029 | 100.00 |
| ELITE-RSLC037842 | 319_GGP | Unmod. | 81.4029 | 81.4029 | 100.00 |
| ELITE-RSLC037842 | 325_GVP | Unmod. | 66.5101 | 66.5101 | 100.00 |
| ELITE-RSLC037842 | 330_GPR | Unmod. | 31.9191 | 31.9191 | 100.00 |
| ELITE-RSLC037844 | 40_GSP | Unmod. | 105.4740 | 105.4740 | 100.00 |
| ELITE-RSLC037844 | 43_GEP | Unmod. | 105.4740 | 105.4740 | 100.00 |
| ELITE-RSLC037844 | 49_GLP | Unmod. | 85.9872 | 85.9872 | 100.00 |
| ELITE-RSLC037844 | 63_GPA | Mod. | 18.1163 | 18.1163 | 100.00 |
| ELITE-RSLC037844 | 66_GPN | Unmod. | 34.2999 | 34.2999 | 100.00 |
| ELITE-RSLC037844 | 70_GIP | Mod. | 18.1163 | 51.7510 | 35.01 |
| ELITE-RSLC037844 | 70_GIP | Unmod. | 33.6348 | 51.7510 | 64.99 |
| ELITE-RSLC037844 | 75_GPA | Unmod. | 35.1084 | 35.1084 | 100.00 |
| ELITE-RSLC037844 | 94_GEP | Unmod. | 37.5798 | 37.5798 | 100.00 |
| ELITE-RSLC037844 | 100_GVP | Mod. | 19.5219 | 19.5219 | 100.00 |
| ELITE-RSLC037844 | 103_GGP | Mod. | 19.5219 | 19.5219 | 100.00 |
| ELITE-RSLC037844 | 145_GQP | Mod. | 19.4177 | 44.3263 | 43.81 |
| ELITE-RSLC037844 | 145_GQP | Unmod. | 24.9086 | 44.3263 | 56.19 |
| ELITE-RSLC037844 | 151_GFP | Unmod. | 40.6166 | 40.6166 | 100.00 |
| ELITE-RSLC037844 | 153_GPK | Mod. | 19.4177 | 62.7464 | 30.95 |
| ELITE-RSLC037844 | 153_GPK | Unmod. | 43.3287 | 62.7464 | 69.05 |
| ELITE-RSLC037844 | 169_GPG | Unmod. | 16.1934 | 16.1934 | 100.00 |
| ELITE-RSLC037844 | 172_GPG | Unmod. | 16.1934 | 16.1934 | 100.00 |
| ELITE-RSLC037844 | 174_GPQ | Unmod. | 16.1934 | 16.1934 | 100.00 |
| ELITE-RSLC037844 | 177_GPP | Unmod. | 16.1934 | 16.1934 | 100.00 |
| ELITE-RSLC037844 | 178_GPP | Unmod. | 16.1934 | 16.1934 | 100.00 |
| ELITE-RSLC037844 | 250_GAP | Mod. | 43.4220 | 43.4220 | 100.00 |
| ELITE-RSLC037844 | 255_GPP | Unmod. | 40.4483 | 40.4483 | 100.00 |
| ELITE-RSLC037844 | 256_GPP | Unmod. | 86.8307 | 86.8307 | 100.00 |
| ELITE-RSLC037844 | 262_GAP | Mod. | 43.1026 | 83.8703 | 51.39 |
| ELITE-RSLC037844 | 262_GAP | Unmod. | 40.7677 | 83.8703 | 48.61 |
| ELITE-RSLC037844 | 316_GEP | Unmod. | 33.4864 | 33.4864 | 100.00 |
| ELITE-RSLC037844 | 319_GGP | Unmod. | 33.4864 | 33.4864 | 100.00 |
| ELITE-RSLC037844 | 325_GVP | Unmod. | 33.4864 | 33.4864 | 100.00 |
| ELITE-RSLC037846 | 40_GSP | Mod. | 19.0729 | 126.0361 | 15.13 |
| ELITE-RSLC037846 | 40_GSP | Unmod. | 106.9632 | 126.0361 | 84.87 |
| ELITE-RSLC037846 | 43_GEP | Mod. | 40.7095 | 177.4290 | 22.94 |
| ELITE-RSLC037846 | 43_GEP | Unmod. | 136.7195 | 177.4290 | 77.06 |
| ELITE-RSLC037846 | 49_GLP | Unmod. | 177.4290 | 177.4290 | 100.00 |
| ELITE-RSLC037846 | 63_GPA | Mod. | 57.5090 | 130.3027 | 44.13 |
| ELITE-RSLC037846 | 63_GPA | Unmod. | 72.7938 | 130.3027 | 55.87 |
| ELITE-RSLC037846 | 66_GPN | Mod. | 54.8535 | 164.7228 | 33.30 |
| ELITE-RSLC037846 | 66_GPN | Unmod. | 109.8693 | 164.7228 | 66.70 |
| ELITE-RSLC037846 | 70_GIP | Mod. | 41.4313 | 115.0897 | 36.00 |
| ELITE-RSLC037846 | 70_GIP | Unmod. | 73.6584 | 115.0897 | 64.00 |
| ELITE-RSLC037846 | 75_GPA | Mod. | 41.4313 | 110.7541 | 37.41 |
| ELITE-RSLC037846 | 75_GPA | Unmod. | 69.3228 | 110.7541 | 62.59 |
| ELITE-RSLC037846 | 94_GEP | Unmod. | 18.3197 | 18.3197 | 100.00 |
| ELITE-RSLC037846 | 103_GGP | Unmod. | 18.3197 | 18.3197 | 100.00 |
| ELITE-RSLC037846 | 145_GQP | Mod. | 43.6728 | 94.3200 | 46.30 |
| ELITE-RSLC037846 | 145_GQP | Unmod. | 50.6472 | 94.3200 | 53.70 |
| ELITE-RSLC037846 | 151_GFP | Mod. | 20.9698 | 20.9698 | 100.00 |
| ELITE-RSLC037846 | 153_GPK | Mod. | 20.9698 | 115.2898 | 18.19 |
| ELITE-RSLC037846 | 153_GPK | Unmod. | 94.3200 | 115.2898 | 81.81 |
| ELITE-RSLC037846 | 169_GPG | Unmod. | 17.8969 | 17.8969 | 100.00 |
| ELITE-RSLC037846 | 172_GPG | Unmod. | 17.8969 | 17.8969 | 100.00 |
| ELITE-RSLC037846 | 174_GPQ | Unmod. | 17.8969 | 17.8969 | 100.00 |
| ELITE-RSLC037846 | 177_GPP | Unmod. | 17.8969 | 17.8969 | 100.00 |
| ELITE-RSLC037846 | 178_GPP | Unmod. | 17.8969 | 17.8969 | 100.00 |
| ELITE-RSLC037846 | 238_GAP | Unmod. | 15.7655 | 15.7655 | 100.00 |
| ELITE-RSLC037846 | 241_GAP | Mod. | 15.7655 | 31.5955 | 49.90 |
| ELITE-RSLC037846 | 241_GAP | Unmod. | 15.8301 | 31.5955 | 50.10 |
| ELITE-RSLC037846 | 250_GAP | Mod. | 15.7655 | 38.0657 | 41.42 |
| ELITE-RSLC037846 | 250_GAP | Unmod. | 22.3003 | 38.0657 | 58.58 |
| ELITE-RSLC037846 | 255_GPP | Mod. | 22.3003 | 45.4473 | 49.07 |
| ELITE-RSLC037846 | 255_GPP | Unmod. | 23.1470 | 45.4473 | 50.93 |
| ELITE-RSLC037846 | 256_GPP | Mod. | 22.3003 | 87.7776 | 25.41 |
| ELITE-RSLC037846 | 256_GPP | Unmod. | 65.4773 | 87.7776 | 74.59 |
| ELITE-RSLC037846 | 262_GAP | Mod. | 45.4473 | 45.4473 | 100.00 |
| ELITE-RSLC037846 | 282_GPP | Unmod. | 18.6874 | 18.6874 | 100.00 |
| ELITE-RSLC037846 | 283_GPP | Unmod. | 18.6874 | 18.6874 | 100.00 |
| ELITE-RSLC037846 | 285_GPP | Unmod. | 18.6874 | 18.6874 | 100.00 |
| ELITE-RSLC037846 | 286_GPP | Unmod. | 18.6874 | 18.6874 | 100.00 |
| ELITE-RSLC037846 | 298_GMP | Unmod. | 18.6874 | 18.6874 | 100.00 |
| ELITE-RSLC037846 | 316_GEP | Unmod. | 124.4596 | 124.4596 | 100.00 |
| ELITE-RSLC037846 | 319_GGP | Unmod. | 124.4596 | 124.4596 | 100.00 |
| ELITE-RSLC037846 | 325_GVP | Mod. | 32.7884 | 124.4596 | 26.34 |
| ELITE-RSLC037846 | 325_GVP | Unmod. | 91.6712 | 124.4596 | 73.66 |
| ELITE-RSLC037848 | 28_GVP | Unmod. | 41.7938 | 41.7938 | 100.00 |
| ELITE-RSLC037848 | 40_GSP | Unmod. | 228.8501 | 228.8501 | 100.00 |
| ELITE-RSLC037848 | 43_GEP | Mod. | 18.8140 | 250.1121 | 7.52 |
| ELITE-RSLC037848 | 43_GEP | Unmod. | 231.2981 | 250.1121 | 92.48 |
| ELITE-RSLC037848 | 49_GLP | Unmod. | 268.6751 | 268.6751 | 100.00 |
| ELITE-RSLC037848 | 58_GAP | Unmod. | 17.1999 | 17.1999 | 100.00 |
| ELITE-RSLC037848 | 63_GPA | Mod. | 18.1023 | 37.4342 | 48.36 |
| ELITE-RSLC037848 | 63_GPA | Unmod. | 19.3319 | 37.4342 | 51.64 |
| ELITE-RSLC037848 | 66_GPN | Mod. | 37.4342 | 91.2268 | 41.03 |
| ELITE-RSLC037848 | 66_GPN | Unmod. | 53.7926 | 91.2268 | 58.97 |
| ELITE-RSLC037848 | 70_GIP | Mod. | 19.3319 | 126.5739 | 15.27 |
| ELITE-RSLC037848 | 70_GIP | Unmod. | 107.2420 | 126.5739 | 84.73 |
| ELITE-RSLC037848 | 75_GPA | Mod. | 19.3319 | 129.3813 | 14.94 |
| ELITE-RSLC037848 | 75_GPA | Unmod. | 110.0494 | 129.3813 | 85.06 |
| ELITE-RSLC037848 | 103_GGP | Unmod. | 21.3252 | 21.3252 | 100.00 |
| ELITE-RSLC037848 | 145_GQP | Mod. | 40.3388 | 108.1789 | 37.29 |
| ELITE-RSLC037848 | 145_GQP | Unmod. | 67.8401 | 108.1789 | 62.71 |
| ELITE-RSLC037848 | 151_GFP | Unmod. | 57.5944 | 57.5944 | 100.00 |
| ELITE-RSLC037848 | 153_GPK | Unmod. | 119.7812 | 119.7812 | 100.00 |
| ELITE-RSLC037848 | 169_GPG | Unmod. | 31.5013 | 31.5013 | 100.00 |
| ELITE-RSLC037848 | 172_GPG | Unmod. | 31.5013 | 31.5013 | 100.00 |
| ELITE-RSLC037848 | 174_GPQ | Unmod. | 31.5013 | 31.5013 | 100.00 |
| ELITE-RSLC037848 | 177_GPP | Unmod. | 31.5013 | 31.5013 | 100.00 |
| ELITE-RSLC037848 | 178_GPP | Unmod. | 31.5013 | 31.5013 | 100.00 |
| ELITE-RSLC037848 | 250_GAP | Mod. | 20.6807 | 20.6807 | 100.00 |
| ELITE-RSLC037848 | 255_GPP | Unmod. | 64.5775 | 64.5775 | 100.00 |
| ELITE-RSLC037848 | 256_GPP | Mod. | 20.6807 | 115.6182 | 17.89 |
| ELITE-RSLC037848 | 256_GPP | Unmod. | 94.9376 | 115.6182 | 82.11 |
| ELITE-RSLC037848 | 262_GAP | Mod. | 68.8397 | 68.8397 | 100.00 |
| ELITE-RSLC037848 | 282_GPP | Unmod. | 20.0323 | 20.0323 | 100.00 |
| ELITE-RSLC037848 | 283_GPP | Unmod. | 20.0323 | 20.0323 | 100.00 |
| ELITE-RSLC037848 | 285_GPP | Unmod. | 20.0323 | 20.0323 | 100.00 |
| ELITE-RSLC037848 | 286_GPP | Unmod. | 20.0323 | 20.0323 | 100.00 |
| ELITE-RSLC037848 | 292_GTP | Unmod. | 20.0323 | 20.0323 | 100.00 |
| ELITE-RSLC037848 | 298_GMP | Unmod. | 20.0323 | 20.0323 | 100.00 |
| ELITE-RSLC037848 | 307_GSP | Mod. | 35.7435 | 35.7435 | 100.00 |
| ELITE-RSLC037848 | 309_GPK | Mod. | 35.7435 | 35.7435 | 100.00 |
| ELITE-RSLC037848 | 316_GEP | Unmod. | 106.9230 | 106.9230 | 100.00 |
| ELITE-RSLC037848 | 319_GGP | Mod. | 17.9221 | 177.4607 | 10.10 |
| ELITE-RSLC037848 | 319_GGP | Unmod. | 159.5386 | 177.4607 | 89.90 |
| ELITE-RSLC037848 | 325_GVP | Unmod. | 177.4607 | 177.4607 | 100.00 |
| ELITE-RSLC037848 | 330_GPR | Unmod. | 56.7676 | 56.7676 | 100.00 |
| ORBI-RSLC037282 | 40_GSP | Unmod. | 50.9065 | 50.9065 | 100.00 |
| ORBI-RSLC037282 | 43_GEP | Unmod. | 50.9065 | 50.9065 | 100.00 |
| ORBI-RSLC037282 | 49_GLP | Unmod. | 50.9065 | 50.9065 | 100.00 |
| ORBI-RSLC037282 | 151_GFP | Unmod. | 21.3525 | 21.3525 | 100.00 |
| ORBI-RSLC037282 | 153_GPK | Unmod. | 21.3525 | 21.3525 | 100.00 |
| ORBI-RSLC037282 | 169_GPG | Unmod. | 16.0640 | 16.0640 | 100.00 |
| ORBI-RSLC037282 | 172_GPG | Unmod. | 16.0640 | 16.0640 | 100.00 |
| ORBI-RSLC037282 | 174_GPQ | Unmod. | 16.0640 | 16.0640 | 100.00 |
| ORBI-RSLC037282 | 177_GPP | Unmod. | 16.0640 | 16.0640 | 100.00 |
| ORBI-RSLC037282 | 178_GPP | Unmod. | 16.0640 | 16.0640 | 100.00 |
| ORBI-RSLC037282 | 255_GPP | Unmod. | 22.7614 | 22.7614 | 100.00 |
| ORBI-RSLC037282 | 256_GPP | Unmod. | 22.7614 | 22.7614 | 100.00 |
| ORBI-RSLC037282 | 262_GAP | Unmod. | 22.7614 | 22.7614 | 100.00 |
| ORBI-RSLC037282 | 270_GPP | Unmod. | 14.6829 | 14.6829 | 100.00 |
| ORBI-RSLC037282 | 271_GPP | Unmod. | 14.6829 | 14.6829 | 100.00 |
| ORBI-RSLC037282 | 273_GPE | Unmod. | 14.6829 | 14.6829 | 100.00 |
| ORBI-RSLC037282 | 282_GPP | Unmod. | 41.6391 | 41.6391 | 100.00 |
| ORBI-RSLC037282 | 283_GPP | Unmod. | 41.6391 | 41.6391 | 100.00 |
| ORBI-RSLC037282 | 285_GPP | Unmod. | 60.3819 | 60.3819 | 100.00 |
| ORBI-RSLC037282 | 286_GPP | Unmod. | 60.3819 | 60.3819 | 100.00 |
| ORBI-RSLC037282 | 292_GTP | Unmod. | 41.6391 | 41.6391 | 100.00 |
| ORBI-RSLC037282 | 298_GMP | Unmod. | 60.3819 | 60.3819 | 100.00 |

